# Population genomics of the invasive Argentine ant (*Linepithema humile*) – adaptive evolution in introduced supercolonies despite low genetic diversity

**DOI:** 10.1101/2024.10.29.620798

**Authors:** Ida Päkkilä, Jenni Paviala, Jes Søe Pedersen, Heikki Helanterä, Lumi Viljakainen

**Author notes:** author for correspondence, Ida Päkkilä, Lumi Viljakainen.

## Abstract

The Argentine ant (*Linepithema humile*), native to South America, has spread globally over the past 150 years, forming extremely large supercolonies in its introduced range. Despite the recent demographic history, including founder effects, Argentine ants thrive in the introduced range and displace native ant species. We adopted a comprehensive approach to investigate selection signals across the whole genome of this highly invasive species, with a specific focus on introduced supercolonies. We also investigated genome-wide diversity and divergence patterns and compared the results with earlier microsatellite marker data studies. We used pooled whole-genome sequence data from 100 workers from the species’ native range and each of the three invasive supercolonies – European Main, Catalonia, and Chile. Invasive supercolonies harboured low genetic diversity and were genetically highly differentiated. Despite this, we could detect signs of positive selection in their genomes. Some of the selected genes showed parallel adaptive evolution in invasive supercolonies. Furthermore, there were no signs that a social organisation based on unrelated workers had led to impaired adaptability (genetic meltdown). We conclude that introduced Argentine ant supercolonies evolve adaptively, indicating that founder effects, low genetic diversity and supercolony social organisation do not always hamper species adaptability.

## 1. Introduction

Invasive species increasingly threaten biodiversity and ecosystems globally (Mollot et al., 2017; Seebens et al., 2017). In the most harmful situations, they cause local extinctions, permanently changing the nature of colonised areas (Holway et al., 2002; Mack et al., 2000). However, the factors contributing to the success of invasive species are still poorly understood, as are their long-term evolutionary prospects in the new ranges, which makes the possibility of controlling these species and mitigating their detrimental effects challenging (Estoup et al., 2016; Pérez et al., 2006; Sherpa & Després, 2021).

Ants exhibit enormous species richness, inhabit diverse ecological niches, and play a crucial role in many ecosystems (Hölldobler & Wilson, 1990; Moreau et al., 2006; Wilson & Hölldobler, 2005). Many ant species are also among the most harmful invasive species (Holway et al., 2002; Lester & Gruber, 2016; McGlynn, 1999). One of these, the highly invasive Argentine ant (*Linepithema humile*), has spread, by accidental human introduction, from its original habitat in South America to all continents except Antarctica during the past 150 years (Suarez et al., 2001; Wetterer et al., 2006, 2009; Wetterer & Wetterer, 2006). Typical of invasive ant species, the Argentine ant also forms large polygynous (multi-queen) and polydomous (multi-nest) societies, known as supercolonies (Pedersen et al., 2006), in which territorial borders between nests are absent over large areas, allowing individuals to move freely between them. In both the native and introduced ranges, these supercolonies form closed breeding units where mating occurs within the nest, and colonies spread by budding (Helanterä, 2022; Pedersen et al., 2006; Vogel et al., 2009). In the introduced range, Argentine ants dominate competition between species, destroy local ecosystems, and cause species extinctions (Giraud et al., 2002; Holway et al., 2002; Sanders et al., 2001, 2003). The Argentine ants’ supercolonies in the introduced range can be enormous, e.g., the European Main supercolony spans over 6,000 kilometres from the Mediterranean coast in Italy to the Spanish Atlantic coast (AntWeb, 2026; Giraud et al., 2002).

Previous research using microsatellite marker data indicated that introduced Argentine ant supercolonies harbour low genetic diversity and are genetically highly differentiated from each other and from the native range supercolonies (Blight et al., 2012; Brandt et al., 2009; Giraud et al., 2002; Jaquiéry et al., 2005; Tsutsui et al., 2000; Vogel et al., 2010). These introduced supercolonies originate from a small number, in some cases only a dozen, of reproducing individuals (Giraud et al., 2002; Vogel et al., 2010), indicating strong founder effects, i.e., loss of genetic variation. Genetic drift (random fluctuations in allele frequencies, often causing allele fixation or loss) is amplified in these small, newly established founder populations, leading to even greater differentiation between populations (Keller & Passera, 1993; Nei et al., 1975). Low initial diversity, combined with strong drift driven by small effective population sizes, should impede adaptive evolution in the introduced Argentine ant supercolonies. Moreover, the general inefficiency of selection on worker traits in supercolonies with low relatedness, due to multiple egg-laying queens, could increase the accumulation of deleterious mutations and decline in fitness (genetic meltdown) (Helanterä et al., 2009; Linksvayer & Wade, 2009). However, the success and dominance of Argentine ants in the introduced range (Vogel et al., 2010) suggest local adaptation to the new environments. This success in novel ranges despite small effective population sizes and decreased selection efficacy raises an important question about the effects of natural selection on the evolution of these ants.

Our earlier population genetic study, focusing on the evolution of immune genes in 18 Argentine ant supercolonies worldwide, including the invasive supercolonies of this study, indicated that, despite recent demographic history, positive selection has affected the evolution of immune genes in the introduced range (Holmberg et al., 2024). In this study, we adopted a more comprehensive approach to investigate signals of selection across the whole genome. We studied native-range supercolonies and three invasive supercolonies from different parts of the world using pooled whole-genome sequence data from 100 workers from the native range and from each of the three invasive supercolonies. We aimed to investigate: i) the overall genome-wide patterns of selection in invasive Argentine ants, i.e., whether signatures of selection could be detected in their genomes, ii) signatures of parallel adaptive evolution among the three invasive supercolonies, and iii) whether genome-wide data confirm the earlier findings, based on microsatellite marker data, about demographic changes during introductions, supercolonial diversity and divergence.

## 2. Materials and Methods

### 2.1 Sampling

We analysed four samples of Argentine ants. One of the samples represents native range supercolonies, and three samples represent invasive supercolonies, one originating from South America and two from Europe (Table S1) (Wild, 2004). All samples include 100 diploid workers. The sample representing the native range supercolonies was collected in Argentina, while the samples representing the three invasive supercolonies were collected in France, Spain and Chile. The three sampled invasive supercolonies arose from different primary introductions from the native range (Vogel et al., 2010). The two European samples were collected from the two known, separate supercolonies: the European Main supercolony (European Main; establishment recorded 1906) and the Catalonian supercolony (Catalonia; 1916), which live partly next to each other along the Mediterranean coastline (Giraud et al., 2002). The third invasive sample from the Chilean supercolony (Chile; 1965) was sampled from the only recognised supercolony in the country (Vogel et al., 2010). For each of the three samples representing the introduced ranges, ten individuals were collected from each of ten different nests within each supercolony (i.e., 100 individuals per supercolony). In the introduced ranges, ten nests were always close together at a single location. For the native-range sample, ten individuals were collected from ten nests each from a separate supercolony from four separate localities (i.e., a total of 100 individuals per ten supercolonies) to capture the total genetic diversity in the species’ natural range (López et al., 2025) (Table S1).

### 2.2. DNA extraction and sequencing

Genomic DNA was extracted individually from each ant using the DNeasy Blood & Tissue Kit (QIAGEN). Equal amounts of DNA from ten individuals per nest were pooled and genome-amplified using REPLI-g Mini Kit (QIAGEN). Finally, all the genome-amplified sub-samples were pooled so that each final sample represented 100 individuals. The samples were sent to BGI Genomics for library preparation and paired-end sequencing using an Illumina Hiseq™ 2000 sequencer, with a read length of 90 base pairs (bp) and an insert size of 200 bp. The clean reads provided by BGI Genomics were filtered to remove adapters, contaminants, and low-quality sequences.

### 2.3. Data processing

The quality encoding of the sequenced raw reads was converted from Phred 64 to Phred 33 (also known as Sanger), as it is more widely used and required in many software programs. The conversion was done using the Seqtk tool (version 1.3) (Li, 2018). After this, raw reads were filtered using trim-fastq.pl script from the basic pipeline of PoPoolation (version 1.2.2) (Kofler et al., 2011). The reads were filtered using an average minimum base quality of 20 and a minimum read length of 70 bp. The quality of the reads was verified using FastQC (version 0.11.8) analyses after filtering (Andrews, 2010).

Filtered reads were aligned to the reference genome (Genome assembly Lhum_UNIL_v1.0, Pan et al., 2024) using the BWA-MEM algorithm from the BWA software package (version 0.7.17) with default settings (Li & Durbin, 2009). The resulting SAM files were sorted, quality-filtered and converted to more compressed binary versions (BAM files) using the view command of SAMtools (version 1.10) (Li et al., 2009). Only aligned reads with a mapping quality of at least 20 and a SAM flag “properly paired” were saved. Duplicates were removed using Picard’s MarkDuplicates tool (version 2.21.4) (*Picard Toolkit*, 2019). RepeatModeler (version 2.0.7) was used to identify repeats, and RepeatMasker (version 4.1.9) was used to mask the identified repeats in the reference genome (Smit & Hubley, 2015). Degeneracy of sites in coding regions was annotated with degenotate (version 1.2.4) (Mirchandani et al., 2024).

### 2.4. Population genetic analyses

#### 2.4.1. Genomic diversity

The genomic diversity of supercolonies was evaluated for the whole genome, as well as separately for four-fold and zero-fold degenerate sites in coding regions. Two genetic diversity estimators, Tajima’s π (π) and Watterson’s θ (θ_w_), were computed using Grenedalf’s (version 0.6.3) diversity command (Czech et al., 2024). Aligned reads were subsampled to a maximum sequencing depth of 200 using the subsampling method subscale. A minimum sequencing depth of four and a minor allele count of two were required per sample. We also estimated microsatellite marker diversity, expected heterozygosity (H_exp_) and allelic richness (k’) for five variable microsatellite loci obtained from Vogel et al. (2010), applying a similar sample size and design as the genomic samples (see Text S2).

#### 2.4.2. Genomic differentiation

To evaluate the genomic differentiation of supercolonies, we calculated Hudson’s estimator of F_ST_using Grenedalf’s fst command (Czech et al., 2024). A minimum sequencing depth of four and a minor allele count of two were required per sample. We also estimated microsatellite marker differentiation using pairwise F_ST_for five variable microsatellite loci obtained from Vogel et al. (2010), applying a sample size and design similar to those of the genomic samples (see Text S2).

#### 2.4.3. Population structure and adaptive evolution

We used the population genomics software BayPass (version 2.31), which utilises a Bayesian framework to identify SNPs subjected to selection and associated with population-specific covariates (Gautier, 2015; Olazcuaga et al., 2020). The different models of BayPass estimate and account for the neutral covariance structure among population allele frequencies resulting from their shared history. Thus, BayPass characterises the neutral genome-wide background pattern of allele-frequency differentiation generated by demographic history and genetic drift and identifies loci that deviate from this expectation, potentially due to selection.

To evaluate population structure and identify SNPs under selection among invasive supercolonies, the analysis was first run in core model mode with default settings. The differentiation outlier statistics, XtX, consider all analysed populations simultaneously and identify SNPs that show unusually high differentiation between populations relative to neutral expectations based on population structure and drift, and thus are potential targets of selection in one or more populations. We analysed the three invasive supercolonies to identify differentiated SNPs in at least one of them, making these SNPs candidates for selection. As this analysis is based on differences in allele frequencies and includes solely invasive supercolonies, it only identifies SNPs that have experienced different selective pressures in at least one of the studied supercolonies, rather than SNPs that have a shared selective regime across all three invasive supercolonies (these SNPs were identified with a different method, see below). XtX statistics were calibrated using a simulated neutral dataset, i.e., a pseudo-observed data set (POD). For neutral data simulation, SNPs were thinned into 20 subsets, each comprising approximately 42K SNPs. These thinned SNP sets were used to estimate the covariance matrix, i.e., Ω of population allele frequencies resulting from their demographic history, which were then used to simulate PODs of 50K SNPs. PODs were then run in core model mode, and the 99th percentiles of XtX differentiation outlier statistics were calculated; the average of these percentiles serves as a threshold to identify outlier SNPs from the real data, i.e., candidates for selection.

Furthermore, a contrast model mode, C2 statistic, was run to estimate whether SNPs are associated with invasiveness. The C2 statistic contrasts and compares allele frequencies between groups of populations defined by the binary covariable of interest, testing for a consistent allele frequency difference between groups. Hence, we contrasted the native-range sample with the group of three invasive supercolonies. This analysis identifies SNPs that share similar allele frequencies across invasive supercolonies and simultaneously differ from those in natives. However, note that the association does not require complete similarity in allele frequencies across all invasive supercolonies, two might be enough. The C2 statistics were calibrated using PODs as described above to indicate whether outlier SNPs were associated with invasiveness. Both models accounted for population structure estimated as Ω of population allele frequencies.

Genes (Annotation GCF_040581485.1-RS_2024_08) containing outlier SNPs with high XtX values were considered as potential targets of positive selection in one or more invasive supercolonies. Genes containing outlier SNPs with high C2 values were associated with invasiveness. We performed gene enrichment analysis using the genes identified as potential targets of positive selection as a test set and the annotated genes of the reference genome (Annotation GCF_040581485.1-RS_2024_08) as a reference set, using the ShinyGO tool (version 0.85) (Ge et al., 2020). Similarly, genes associated with invasiveness were analysed for enrichment. Moreover, genes identified by both the XtX and C2 statistics were analysed for enrichment.

## 3. Results

### 3.1. Properties of the data

Sequencing reads covered 92.78%, 93.11%, 82.71%, and 92.38% of the reference genome in the native-range sample, Main, Catalonia and Chile, while the mean sequencing depths were 87, 98, 61, and 78, respectively. Repetitive regions, which comprised 31% of the reference genome, were removed from all downstream analyses.

### 3.2. Introduced supercolonies are expanding and harbour low genetic diversity

We evaluated the genetic diversity of supercolonies with two genetic diversity estimators, Tajima’s pairwise nucleotide diversity (π) and Watterson’s theta (θ_w_), describing the population mutation rate. To ensure comparability, we accepted only sites with data from all supercolonies for diversity calculations. These sites contained 502,659 whole-genome SNPs, of which 59,575 were 4-fold SNPs, and 122,678 were 0-fold SNPs. SNPs were identified simultaneously across the entire data set.

According to π, the native-range supercolonies were more genetically diverse than the invasive supercolonies (p-values < 0.05 in all pairwise comparisons, see details in Table S3; Table 1). Both expected heterozygosity (H_exp_) and allelic richness (k’) estimates derived from microsatellite marker diversity yielded similar results to π – the sample of native-range supercolonies harbours higher diversity than the invasive supercolonies (Table S4, Figure S5). Instead, θ_w_ indicated that Catalonia had the highest genetic diversity (p-values < 0.05 in all pairwise comparisons, see details in Table S3; Table 1). The Catalonia supercolony harboured an excess of rare alleles, as evidenced also by the high peaks on the left side of the folded site frequency spectrum (Figure 1). For each supercolony, θ_w_ was always higher than π, especially pronounced in the invasive supercolonies, indicating expanding population sizes (p-values < 0.05 in all pairwise comparisons, see details in Table S3; Table 1). Moreover, four-fold sites had consistently higher diversity than zero-fold sites, as expected because synonymous four-fold sites are generally subjected to weaker purifying selection than nonsynonymous zero-fold sites (p-values < 0.05 in all pairwise comparisons, see details in Table S3; Table 1).

**Table 1.**
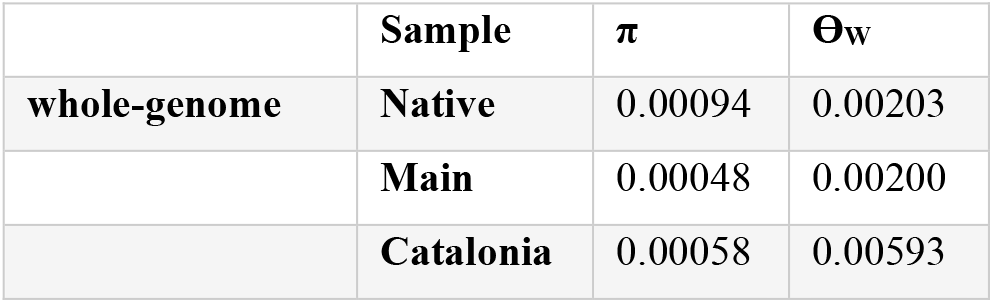
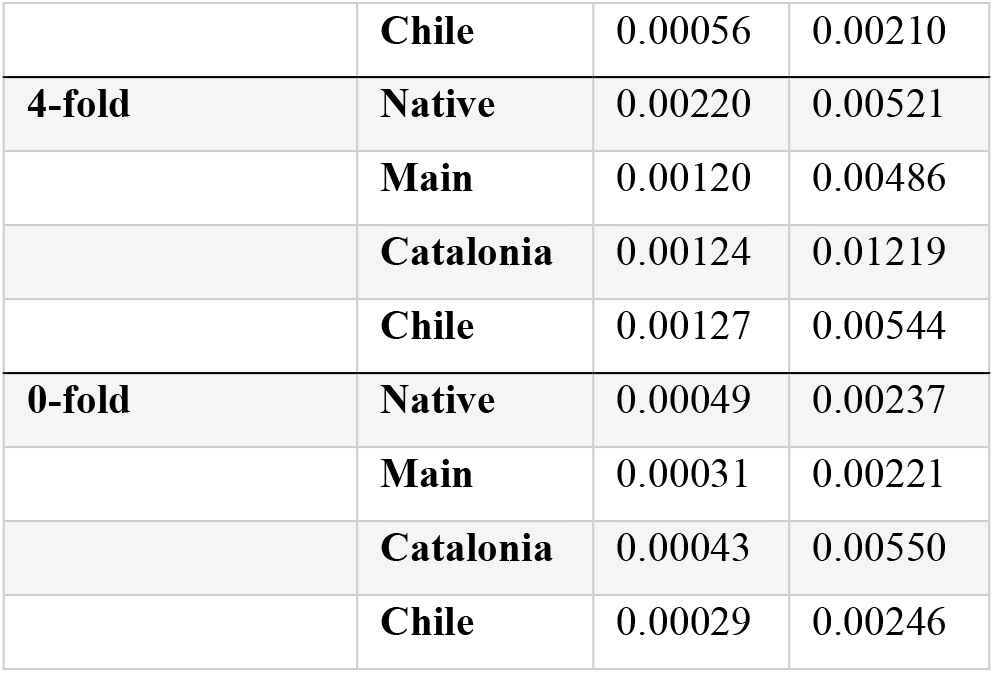
The average per-site diversity estimators, π and θ_W_, for whole-genome, four-fold sites, and zero-fold sites.

| | Sample | $\pi$ | $\Theta_w$ |
| --- | --- | --- | --- |
| whole-genome | Native | 0.00094 | 0.00203 |
|  | Main | 0.00048 | 0.00200 |
|  | Catalonia | 0.00058 | 0.00593 |
|  | **Chile** | 0.00056 | 0.00210 |
| **4-fold** | **Native** | 0.00220 | 0.00521 |
|  | **Main** | 0.00120 | 0.00486 |
|  | **Catalonia** | 0.00124 | 0.01219 |
|  | **Chile** | 0.00127 | 0.00544 |
| **0-fold** | **Native** | 0.00049 | 0.00237 |
|  | **Main** | 0.00031 | 0.00221 |
|  | **Catalonia** | 0.00043 | 0.00550 |
|  | **Chile** | 0.00029 | 0.00246 |

**Figure 1.**
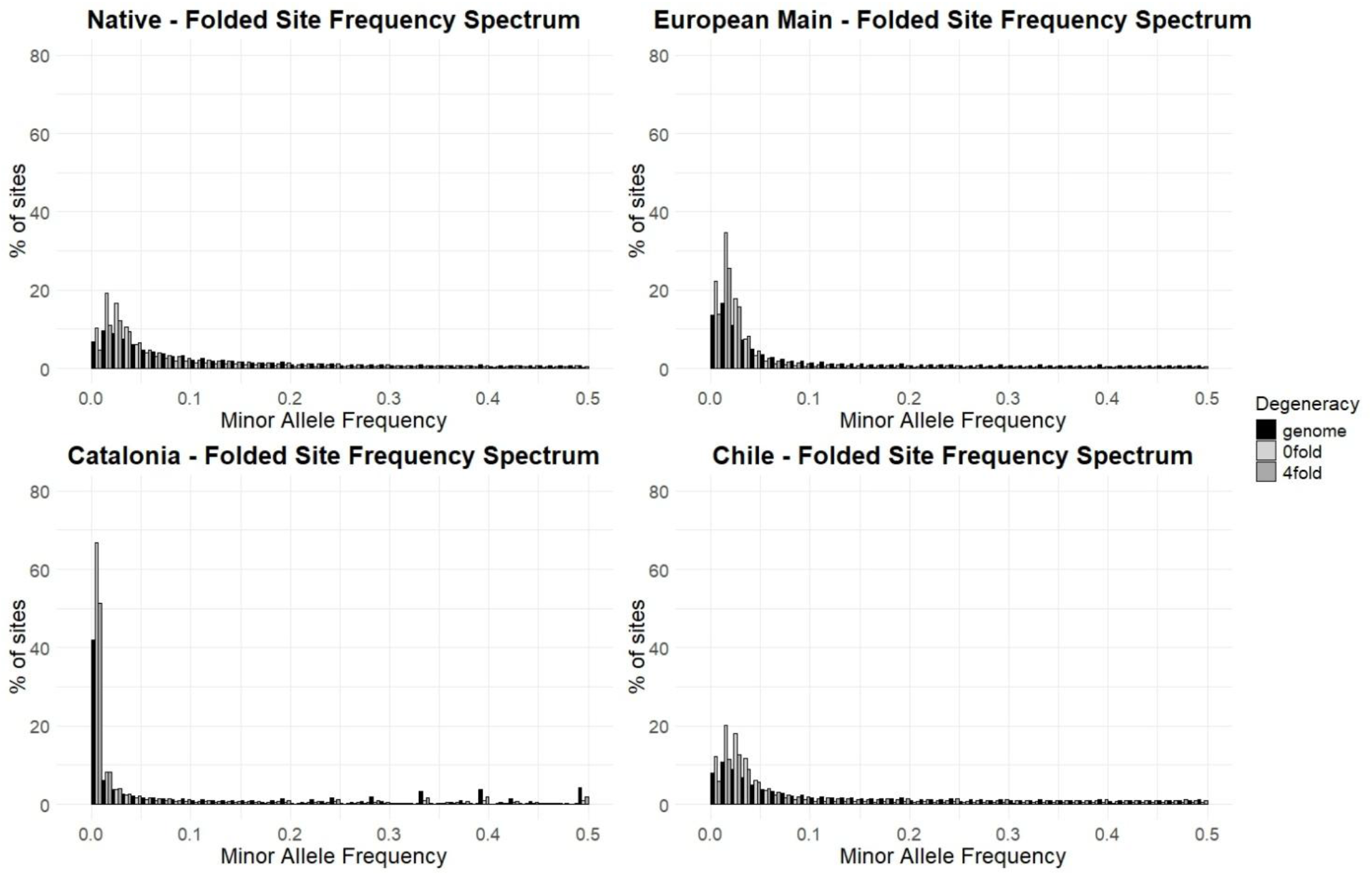
Folded site frequency spectra for Native, Main, Catalonia, and Chile.

### 3.3. Introduced supercolonies are genetically different from each other

We used pairwise F_ST_ values to evaluate genetic differentiation between supercolonies. The results indicated that supercolonies were genetically highly differentiated from each other (Table 2). The pairwise F_ST_ values were consistently highest between Catalonia and the other supercolonies, indicating that Catalonia was the most genetically differentiated supercolony (Table 2). The microsatellite marker data also revealed genetic differences between supercolonies (Table S6, Figure S7). The marker data indicated that invasive supercolonies were more differentiated from each other than from the native-range supercolonies (Table S6, Figure S7) – a pattern that we did not observe in the genomic data.

**Table 2.** Genetic differences measured as pairwise F_ST_ values, genome-wide average.

| $F_{ST}$ | Native | Main | Catalonia | Chile |
| --- | --- | --- | --- | --- |
| **Native** | 0 | 0.2812 | 0.4434 | 0.2670 |
| **Main** |  | 0 | 0.5034 | 0.2507 |
| **Catalonia** |  |  | 0 | 0.5148 |
| **Chile** |  |  |  | 0 |

### 3.4. Argentine ants can adapt to new environments

The core model and the XtX statistics of the BayPass software were used to identify candidate genes for positive selection among invasive supercolonies. 1220 genes (see Table S8) contained one or more SNPs with an XtX value greater than the calibrated threshold (6.3825), indicating them as outlier loci in the genome scan for adaptive differentiation, and thus candidates for positive selection in one or more of the invasive supercolonies. The XtX outlier SNPs were distributed across the genome (Figure S9). Enriched molecular function and biological process GO terms among the candidate genes were mainly related to nervous system development, ion transport, signalling-related processes, and transcriptional regulation (Table S10).

A contrast analysis based on C2 statistics of the BayPass software was used to identify genes associated with invasiveness. 285 genes (see Table S11) contained one or more SNPs with a C2 value greater than the calibrated threshold (4.7973), indicating them as outlier loci in the genome scan for association with invasiveness. Thus, these genes showed signs of parallel adaptive evolution across invasive supercolonies. These C2 outlier SNPs were also distributed across the genome (Figure S12). Enriched molecular function and biological process GO terms among genes associated with invasiveness were mainly related to ion transport, cellular signalling and communication, and gene regulation and transcriptional processes (Table S13). Notably, 97 genes were identified by both XtX and C2 analyses. These genes showed both high differentiation between invasive supercolonies and an association with invasiveness, indicating parallel adaptive evolution at each gene in two out of the three invasive supercolonies studied. One molecular function GO term, GO:0030125 Semaphorin receptor binding, was enriched among these genes (Table S14). The remaining 188 genes associated with invasiveness (high C2) were those that showed parallel adaptive evolution across all three invasive supercolonies (as not identified with the XtX).

Both models estimated and accounted for the covariance structure among population allele frequencies resulting from their shared demographic history. The SVD-based visualisation of the Ω matrix summarises the genetic structure among the native-range sample and the three invasive supercolonies (Figure 2). PC1 explains approximately 50% of the total genetic variation and distinguishes native-range supercolonies from invasive ones. PC2 explains approximately 35% of the variation and distinguishes the Catalonia supercolony from the other invasive supercolonies (Figure 2).

**Figure 2.**
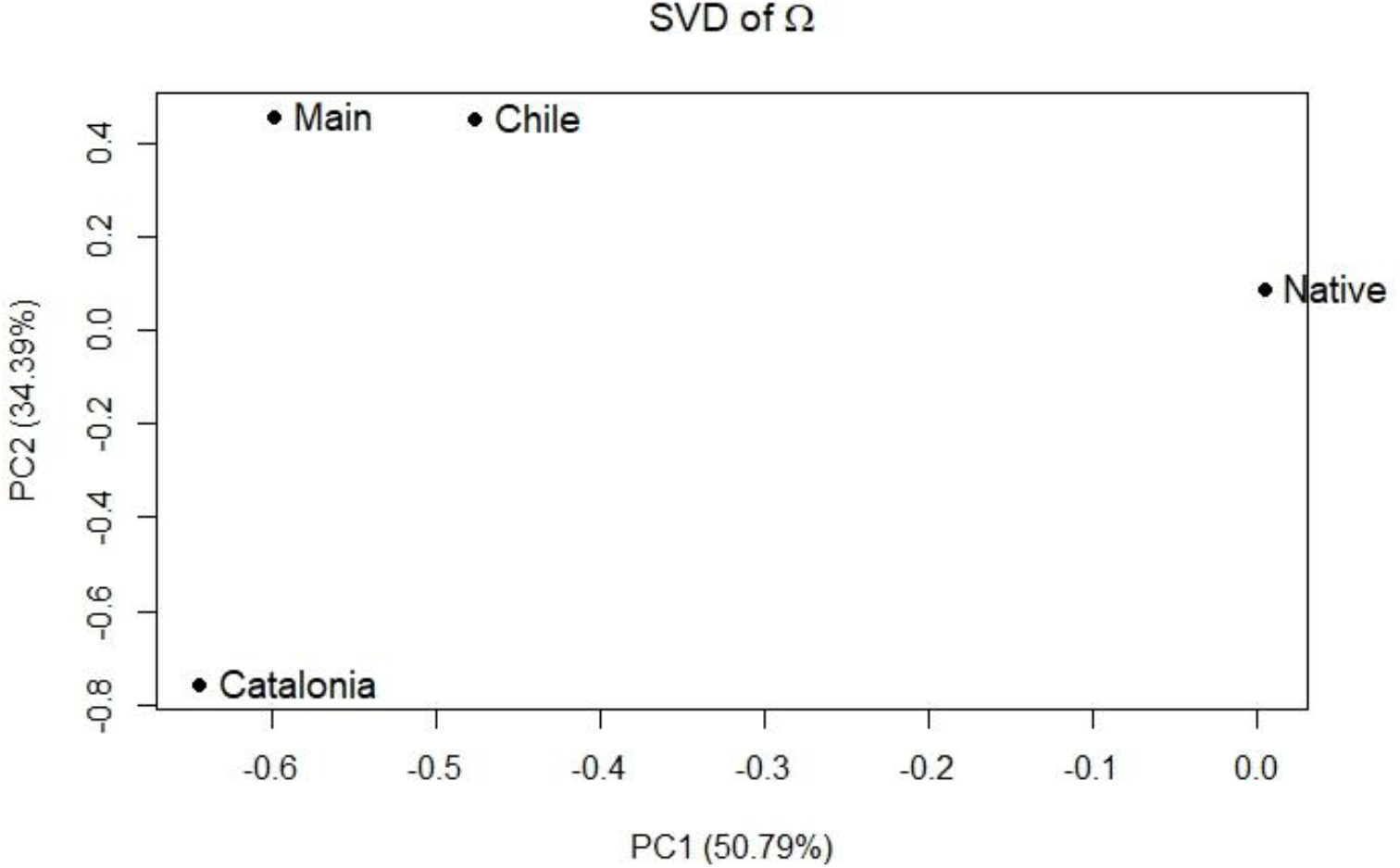
The SVD-based visualisation of the covariance matrix, i.e., Ω of population allele frequencies.

## 4. Discussion

This study aimed to shed light on the highly invasive Argentine ant’s recent evolutionary history and genome dynamics, specifically focusing on the introduced supercolonies. We searched for answers to the following questions: i) can signs of selection be detected in the genomes of the highly invasive Argentine ant?, ii) are there signatures of parallel adaptive evolution among invasive supercolonies?, iii) are we able to verify, using genomic data, the earlier population genetic findings on diversity and divergence based on microsatellite marker data?

We detected signs of selection in the invasive supercolonies, indicating adaptability. Importantly, the method we used accounts for the neutral demographic processes. This is particularly important for recently established invasive supercolonies, where founder effects and subsequent genetic drift are expected to have strongly shaped neutral genetic variation and may otherwise generate signals that mimic selection. Numerous genes were identified as candidates for positive selection among invasive supercolonies. Due to the global nature of the analysis considering allele frequency differences, it cannot be said whether these genes were putatively positively selected in one or more of the supercolonies, or, if selected in more than one, whether the selected allele is the same or different. But it is evident that positive selection shapes the evolution of invasive Argentine ant supercolonies. Moreover, some genes were associated with invasiveness, indicating parallel adaptive evolution in invasive supercolonies. Thus, selective regimes are probably not entirely different between invasive supercolonies, and some selective pressures are shared across invasive ranges. Whilst the Mediterranean-like climate is shared across the native and invasive ranges of the Argentine ant (Roura-Pascual et al., 2011), other biotic and abiotic factors, such as pathogens and competitors, likely differ between regions, leading to distinct local selective pressures and local adaptation (Holmberg et al., 2024; Lester & Gruber, 2016). As the native range supercolonies were analysed as a single group rather than separately, our approach provides a broad overview of genetic variation across the native range. Thus, if native range supercolonies’ allele frequencies differed, this would have led to averaged allele frequencies of the native-range sample and reduced power to detect some differentiation signals between allele frequencies of native-range sample and invasive supercolonies (high XtX values). Nevertheless, the differentiation signals identified and reported here between native-range sample and invasive supercolonies were sufficiently strong to remain detectable despite the native range sampling strategy, suggesting that our results are conservative.

We detected candidates for positive selection in several immune genes (emp, imd, hop, bsk, Trax, Toll-6, and Sp1) in the whole-genome scanning. We could not find overlap between the positively selected immune genes identified here and those reported in the earlier Argentine ant immune gene study, even though the studied supercolonies are the same (worker specimens differ) (Holmberg et al., 2024). This might be due to differences in sampling strategies and the methods used. However, together, these findings indicate that immune genes play important roles in how invasive Argentine ants adapt to new ranges.

Wagner et al. (2023) reported that Argentine ants were excellent olfactory learners. They proposed that efficient learning combined with other features of Argentine ants, such as rapid discovery of food sources and other resources, and rapid recruitment, might provide a competitive advantage over local species in introduced ranges (Wagner et al., 2023). Intriguingly, the candidate genes for positive selection include olfactory learning (trp, mura, mnb, Mob2, cher, PKA-R1, Fas3, Fas1) and memory genes (eag, aPKC, dnc).

The enriched GO terms among the candidate genes for positive selection included a range of molecular functions. Some of these overlap with GO terms for genes that have evolved under positive selection across the ant phylogeny over a long evolutionary time (Barkdull & Moreau, 2023). These GO terms were GO:0015103 Inorganic anion transmembrane transporter activity, GO:0022836 Gated channel activity, GO:0022857 Transmembrane transporter activity, GO:0004930 G protein-coupled receptor activity, GO:0004672 Protein kinase activity, GO:0005515 Protein binding, and GO:0008061 Chitin binding.

The general observation of low diversity and high divergence in the invasive supercolonies based on our genome-wide data largely confirms the results of earlier marker data studies of Argentine ants (Blight et al., 2012; Brandt et al., 2009; Giraud et al., 2002; Jaquiéry et al., 2005; Tsutsui et al., 2000; Vogel et al., 2010). However, there were some discrepancies between the results of genomic data and the microsatellite loci obtained from Vogel et al. (2010). Catalonia appeared to be even more differentiated from other supercolonies based on the genomic data than the marker data suggested, with an excess of rare alleles (high θ_w_). Similarly, in another invasive ant species, *Cardiocondyla obscurior*, genome-wide evidence showed reduced genetic diversity in invasive populations (Errbii et al., 2021). Moreover, marker data have demonstrated reduced genetic diversity in several other invasive ant populations of *Nylanderia fulva, Pheidole megacephala, Solenopsis invicta, Solenopsis geminata*, and *Wasmannia auropunctata* (Fournier et al., 2005, 2009, 2012; Ross et al., 1993, 1996; Wauters et al., 2018). Together, these results indicate that introduced populations of invasive ant species typically harbour less genetic diversity than native populations. This is in line with the typical supercolonial lifestyle of the invasive ants, i.e., the formation of closed breeding units and the fact that populations in the introduced ranges usually comprise single supercolonies.

Our results give no indication that the demographic and ecological changes that accompany an invasion, namely the expansion of supercolonies in the absence of competition that drastically reduces the relatedness between sterile workers and reproductive queens (Giraud et al., 2002; Helanterä, 2022; Helanterä et al., 2009; Pedersen et al., 2006), would have amplified the predicted accumulation of harmful mutations or hampered positive selection on worker traits (Linksvayer & Wade, 2009) in the invasive supercolonies. This could be due to, firstly, the relatively short time span – less than 100 generations – since introductions, or secondly, the rarity of genes only expressed in workers, where such mutation accumulation is predicted and shown to occur (Linksvayer & Wade, 2009; Warner et al., 2017). The rarity of such genes could mask any genome-wide effect, and testing this hypothesis in invasive supercolonies requires high-quality gene expression data.

The high divergence between the invasive Argentine ant supercolonies included in this study may be explained by the fact that they have originated from different primary introductions from the native range without secondary reinforcements (Giraud et al., 2002; Vogel et al., 2010). Moreover, founder effect likely initiated the first shift toward genetic differentiation within the introduced supercolonies, as the effective queen number establishing the European Main as well as the Catalonian supercolonies has been estimated to be between 6 to 13 individuals (Giraud et al., 2002; Vogel et al., 2010), while in the native range supercolonies, it has been estimated to be extremely high, reaching infinity (Pedersen et al., 2006). Subsequent gene flow between invasive supercolonies has been prevented by two factors: i) Argentine ant workers are highly aggressive toward males originating from other colonies, and ii) there is considerable geographic distance between the studied supercolonies, except the European Main and Catalonian supercolonies, which live partly next to each other (Blight et al., 2012; Giraud et al., 2002; Jaquiéry et al., 2005; Vogel et al., 2010). Although the European Main and Catalonia supercolonies live close to each other, there is no gene flow between them (Blight et al., 2012; Giraud et al., 2002; Jaquiéry et al., 2005). In addition, amplified genetic drift in small founder populations might have further increased fluctuations in allele frequencies, contributing to diversification within expanding supercolonies (Nei et al., 1975; Vogel et al., 2010). Especially in Catalonia, the excess of rare alleles might indicate both substantial expansion after population foundation and strong purifying selection. Other social insects, such as the yellow-legged hornet *Vespa velutina* and German wasp *Vespula germanica*, have also successfully invaded regions outside their native range despite low genetic diversity (Arca et al., 2015; Brenton-Rule et al., 2018; Eloff et al., 2020; Takeuchi et al., 2017). In the yellow-legged hornet, successful invasion possibly originated from as few individuals as just one (one queen mated with multiple males) (Arca et al., 2015; Takeuchi et al., 2017). In addition, other non-social insects, such as the African fig fly *Zaprionus indianus* (Comeault et al., 2020), and other species, such as the invasive wart comb jelly *Mnemiopsis leidyi* (Jaspers et al., 2021) have also proven that invasion success does not require high genomic diversity. Thus, the general assumption of invasive populations’ lower adaptive potential is incorrect, as the founder effect and low diversity do not always prevent invaders’ success.

The success of a wide range of invasive species populations with reduced adaptive potential due to low genetic diversity has been explained by factors such as gene flow, stress-induced accelerated mutation rate or TE-activation, epigenetic changes, pre-adaptation in the native range, and the release from natural pathogens and enemies (Errbii et al., 2021; Estoup et al., 2016; Manfredini et al., 2019; Marin et al., 2019; North et al., 2021; Schrader et al., 2014). Despite the low genetic diversity, the invasive Argentine ants continue to evolve adaptively. This indicates that founder effects, low genetic diversity and supercolony social organisation do not always hamper species adaptability. Future comparative population genomic studies incorporating a broader sampling of both native and introduced supercolonies, together with high-quality genomic and transcriptomic data at the individual level, would provide an important opportunity to further evaluate the detected signals of selection and determine how consistently they are replicated across independently introduced populations, i.e., the extent of parallel adaptive evolution among invasive supercolonies. Furthermore, the precise functional roles of the candidate genes identified as targets of positive selection remain to be elucidated within the complex biological pathways and processes underlying invasion success. By identifying genomic regions potentially associated with adaptation in invasive populations, our study provides an excellent groundwork and frame for future studies of the Argentine ant in the field of invasion genomics.

