## Supplemental file for "Population genomics of the invasive Argentine ant (*Linepithema humile*) – adaptive evolution in introduced supercolonies despite low genetic diversity" (from another version of this preprint: version 2, 2026-04-18)

**Table of Contents:**


| |
|---|
| **Table S1: Sampling locations** |
| **Text S2: Comparing the results of genomic data and microsatellite marker data** |
| **Table S3: Results of statistical tests** |
| **Table S4: Microsatellite marker diversity ($H_{exp}$ & k')** |
| **Figure S5: Genomic diversity (π & $\Theta_w$) versus microsatellite marker diversity ($H_{exp}$ & k')** |
| **Table S6: Microsatellite marker differentiation ($F_{ST}$)** |
| **Figure S7: Genomic differentiation ($F_{ST}$) versus microsatellite marker differentiation ($F_{ST}$)** |
| **Table S8: Positively selected genes based on the XtX statistic** |
| **Figure S9. Positively selected SNPs based on the XtX statistic** |
| **Table S10. Enriched GO terms among positively selected genes** |
| **Table S11. Genes associated with invasiveness based on the C2 statistics** |
| **Figure S12. SNPs associated with invasiveness based on the C2 statistics** |
| **Table S13. Enriched GO terms among genes associated with invasiveness** |
| **Table S14. Enriched GO terms among genes identified with XtX and C2 (overlap)** |


**Table S1.** Sampling locations of the Argentine ant supercolonies studied. Sampling was conducted in four geographic areas to represent both the native and introduced ranges. In the native range, ten individuals were collected from the ten different nests belonging to ten different supercolonies (a total of 100 individuals) from four different localities in the native range. These 100 individuals from the native range were pooled to represent the final native-range sample. In the introduced range, ten individuals were collected from the ten different nests belonging to the same supercolony (a total of 100 individuals) in each of the three introduced ranges. In the case of invasive supercolonies, these 100 individuals were pooled to represent the final invasive supercolony-specific samples – The European Main, The Catalonian, and The Chilean supercolonies. The native-range sample captures the full genetic diversity within the species' native range, rather than representing a single native supercolony. Sampled nests were a subset of those used in Vogel et al. (2010), but the worker specimens differed.

| Supercolony name | Population genomic sample name | Geographic range [native (N) or introduced (I)] | Locality | Supercolony ID | Nest sample ID [ten workers from each nest] |
|---|---|---|---|---|---|
| The native supercolonies | Native | N | Argentina, Corrientes, Arroyo Cuay Grande | Corrientes S1 | 14 |
| The native supercolonies | Native | N | Argentina, Corrientes, Arroyo Cuay Grande | Corrientes S2 | 21 |
| The native supercolonies | Native | N | Argentina, Buenos Aires, Otamendi | Otamendi S2 | 106 |

| | | | | | |
|---|---|---|---|---|---|
| The native supercolonies | Native | N | Argentina, Buenos Aires, Otamendi | Otamendi S3 | 70 |
| The native supercolonies | Native | N | Argentina, Buenos Aires, Otamendi | Otamendi S4 | 65 |
| The native supercolonies | Native | N | Argentina, Buenos Aires, Otamendi | Otamendi S6 | 30 |
| The native supercolonies | Native | N | Argentina, Buenos Aires, Otamendi | Otamendi S9 | 01 |
| The native supercolonies | Native | N | Argentina, Buenos Aires, Buenos Aires City, Boca | Boca S1 | 02 |
| The native supercolonies | Native | N | Argentina, Buenos Aires, Buenos Aires City, Boca | Boca S2 | 07 |
| The native supercolonies | Native | N | Argentina, Buenos Aires, Santa Coloma | Santa Coloma | 04 |
| The European Main supercolony | European Main | I | France, Port-Leucate | European Main | 58, 60, 61, 63, 64, 65, 68, 71, 76, 80 |
| The Catalonian supercolony | Catalonia | I | Spain, Pals | Catalonia | 03, 14, 15, 17, 18, 20, 22, 24, 25, 33 |
| The Chilean supercolony | Chile | I | Chile, Santiago | Chile | 02, 04, 05, 06, 07, 08, 09, 10, 11, 13 |

Vogel, V., Pedersen, J. S., Giraud, T., Krieger, M. J. B., & Keller, L. (2010). The worldwide expansion of the Argentine ant. Diversity and Distributions, 16(1), 170–186. https://doi.org/10.1111/j.1472-4642.2009.00630.x

**Text S2.** Comparing the results of genomic data and microsatellite marker data.

Material and methods: Microsatellite marker data on five variable microsatellite loci (Lhum-3, Lhum-11, Lhum-19, Lhum-28, and Lhum-35) were obtained from Vogel et al. (2010), applying a similar sample size and design as the genomic samples, i.e. native with the same ten supercolonies (ten worker individuals each) as subsamples and each of the three introduced supercolonies with ten nests as subsamples. Estimations of genetic diversity (expected heterozygosity, $H_{exp}$, and allelic richness, k') and genetic differentiation (pairwise $F_{ST}$) were performed as in Vogel et al. (2010) with dispersal measures (SE) calculated by jackknifing over subsamples.

Vogel, V., Pedersen, J. S., Giraud, T., Krieger, M. J. B., & Keller, L. (2010). The worldwide expansion of the Argentine ant. Diversity and Distributions, 16(1), 170–186. https://doi.org/10.1111/j.1472-4642.2009.00630.x

**Table S3.** Results of statistical tests.

| Data | Comparison group | Tested parameter | Statistical test | Test statistic | Sample size | P-value |
|---|---|---|---|---|---|---|
| Whole-genome SNPs | Native vs. Main | π | Wilcoxon test | 8.8468E+10 | $n_{Native}$=502659, $n_{Main}$=502659 | <2.2e$^{-16}$ |
| Whole-genome | Native vs. Catalonia | π | Wilcoxon test | 7.2327E+10 | $n_{Native}$=502659, $n_{Catalonia}$=502659 | <2.2e$^{-16}$ |

| | | | | | | |
|---|---|---|---|---|---|---|
| SNPs | | | | | | |
| Whole-genome SNPs | Native vs. Chile | π | Wilcoxon test | 9.0262E+10 | $n_{Native}$=502659, $n_{Chile}$=502659 | <2.2e$^{-16}$ |
| Whole-genome SNPs | Main vs. Catalonia | π | Wilcoxon test | 1.4765E+11 | $n_{Main}$=502659, $n_{Catalonia}$=502659 | <2.2e$^{-16}$ |
| Whole-genome SNPs | Main vs. Chile | π | Wilcoxon test | 1.2691E+11 | $n_{Main}$=502659, $n_{Chile}$=502659 | 3.778e-06 |
| Whole-genome SNPs | Catalonia vs. Chile | π | Wilcoxon test | 1.0700E+11 | $n_{Catalonia}$=502659, $n_{Chile}$=502659 | <2.2e$^{-16}$ |
| Whole-genome SNPs | Native vs. Main | $\Theta_w$ | Wilcoxon test | 1.1927E+11 | $n_{Native}$=502659, $n_{Main}$=502659 | <2.2e$^{-16}$ |
| Whole-genome SNPs | Native vs. Catalonia | $\Theta_w$ | Wilcoxon test | 2.1777E+11 | $n_{Native}$=502659, $n_{Catalonia}$=502659 | <2.2e$^{-16}$ |
| Whole-genome SNPs | Native vs. Chile | $\Theta_w$ | Wilcoxon test | 1.3667E+11 | $n_{Native}$=502659, $n_{Chile}$=502659 | <2.2e$^{-16}$ |
| Whole-genome SNPs | Main vs. Catalonia | $\Theta_w$ | Wilcoxon test | 3.3311E+10 | $n_{Main}$=502659, $n_{Catalonia}$=502659 | <2.2e$^{-16}$ |
| Whole-genome SNPs | Main vs. Chile | $\Theta_w$ | Wilcoxon test | 1.0909E+11 | $n_{Main}$=502659, $n_{Chile}$=502659 | <2.2e$^{-16}$ |
| Whole-genome SNPs | Catalonia vs. Chile | $\Theta_w$ | Wilcoxon test | 2.1489E+11 | $n_{Catalonia}$=502659, $n_{Chile}$=502659 | <2.2e$^{-16}$ |
| Whole-genome SNPs | Native π vs. Native $\Theta_w$ | π & $\Theta_w$ | Wilcoxon test | 4.8191E+10 | $n_{Native(\pi)}$=502659, $n_{Native(\Theta w)}$=502659 | <2.2e$^{-16}$ |
| Whole-genome SNPs | Main π vs. Main $\Theta_w$ | π & $\Theta_w$ | Wilcoxon test | 2.5431E+10 | $n_{Main(\pi)}$=502659, $n_{Main(\Theta w)}$=502659 | <2.2e$^{-16}$ |
| Whole-genome SNPs | Catalonia π vs. Catalonia $\Theta_w$ | π & $\Theta_w$ | Wilcoxon test | 1.2672E+10 | $n_{Catalonia(\pi)}$=502659, $n_{Catalonia(\Theta w)}$=502659 | <2.2e$^{-16}$ |
| Whole-genome SNPs | Chile π vs. Chile $\Theta_w$ | π & $\Theta_w$ | Wilcoxon test | 3.0211E+10 | $n_{Chile(\pi)}$=502659, $n_{Chile(\Theta w)}$=502659 | <2.2e$^{-16}$ |
| 4-fold SNPs | Native vs. Main | π | Wilcoxon test | 1.3811E+09 | $n_{Native}$=59575, $n_{Main}$=59575 | <2.2e$^{-16}$ |
| 4-fold SNPs | Native vs. Catalonia | π | Wilcoxon test | 1.1087E+09 | $n_{Native}$=59575, $n_{Catalonia}$=59575 | <2.2e$^{-16}$ |
| 4-fold SNPs | Native vs. Chile | π | Wilcoxon test | 1.2836E+09 | $n_{Native}$=59575, $n_{Chile}$=59575 | <2.2e$^{-16}$ |
| 4-fold | Main vs. | π | Wilcoxon | 2.1511E+09 | $n_{Main}$=59575, | <2.2e$^{-16}$ |

| | | | | | | |
|---|---|---|---|---|---|---|
| SNPs | Catalonia | | test | | $n_{Catalonia}$=59575 | |
| 4-fold SNPs | Main vs. Chile | π | Wilcoxon test | 1.9404E+09 | $n_{Main}$=59575, $n_{Chile}$=59575 | $<2.2e^{-16}$ |
| 4-fold SNPs | Catalonia vs. Chile | π | Wilcoxon test | 1.5934E+09 | $n_{Catalonia}$=59575, $n_{Chile}$=59575 | $<2.2e^{-16}$ |
| 4-fold SNPs | Native vs. Main | $\Theta_w$ | Wilcoxon test | 1.2072E+09 | $n_{Native}$=59575, $n_{Main}$=59575 | $2.2e^{-16}$ |
| 4-fold SNPs | Native vs. Catalonia | $\Theta_w$ | Wilcoxon test | 2.7668E+09 | $n_{Native}$=59575, $n_{Catalonia}$=59575 | $2.2e^{-16}$ |
| 4-fold SNPs | Native vs. Chile | $\Theta_w$ | Wilcoxon test | 2.0709E+09 | $n_{Native}$=59575, $n_{Chile}$=59575 | $2.2e^{-16}$ |
| 4-fold SNPs | Main vs. Catalonia | $\Theta_w$ | Wilcoxon test | 6.3327E+08 | $n_{Main}$=59575, $n_{Catalonia}$=59575 | $2.2e^{-16}$ |
| 4-fold SNPs | Main vs. Chile | $\Theta_w$ | Wilcoxon test | 9.6040E+08 | $n_{Main}$=59575, $n_{Chile}$=59575 | $2.2e^{-16}$ |
| 4-fold SNPs | Catalonia vs. Chile | $\Theta_w$ | Wilcoxon test | 2.6810E+09 | $n_{Catalonia}$=59575, $n_{Chile}$=59575 | $2.2e^{-16}$ |
| 4-fold SNPs | Native π vs. Native $\Theta_w$ | π & $\Theta_w$ | Wilcoxon test | 6.0410E+08 | $n_{Native(\pi)}$=59575, $n_{Native(\Theta w)}$=59575 | $<2.2e^{-16}$ |
| 4-fold SNPs | Main π vs. Main $\Theta_w$ | π & $\Theta_w$ | Wilcoxon test | 3.3156E+08 | $n_{Main(\pi)}$=59575, $n_{Main(\Theta w)}$=59575 | $<2.2e^{-16}$ |
| 4-fold SNPs | Catalonia π vs. Catalonia $\Theta_w$ | π & $\Theta_w$ | Wilcoxon test | 1.8695E+08 | $n_{Catalonia(\pi)}$=59575, $n_{Catalonia(\Theta w)}$=59575 | $<2.2e^{-16}$ |
| 4-fold SNPs | Chile π vs. Chile $\Theta_w$ | π & $\Theta_w$ | Wilcoxon test | 3.7326E+08 | $n_{Chile(\pi)}$=59575, $n_{Chile(\Theta w)}$=59575 | $<2.2e^{-16}$ |
| 0-fold SNPs | Native vs. Main | π | Wilcoxon test | 7.3222E+09 | $n_{Native}$=122678, $n_{Main}$=122678 | $2.2e^{-16}$ |
| 0-fold SNPs | Native vs. Catalonia | π | Wilcoxon test | 5.8672E+09 | $n_{Native}$=122678, $n_{Catalonia}$=122678 | $2.2e^{-16}$ |
| 0-fold SNPs | Native vs. Chile | π | Wilcoxon test | 6.3093E+09 | $n_{Native}$=122678, $n_{Chile}$=122678 | $2.2e^{-16}$ |
| 0-fold SNPs | Main vs. Catalonia | π | Wilcoxon test | 9.2926E+09 | $n_{Main}$=122678, $n_{Catalonia}$=122678 | $2.2e^{-16}$ |
| 0-fold SNPs | Main vs. Chile | π | Wilcoxon test | 8.7092E+09 | $n_{Main}$=122678, $n_{Chile}$=122678 | $2.2e^{-16}$ |
| 0-fold SNPs | Catalonia vs. Chile | π | Wilcoxon test | 7.0275E+09 | $n_{Catalonia}$=122678, $n_{Chile}$=122678 | $2.2e^{-16}$ |
| 0-fold SNPs | Native vs. Main | $\Theta_w$ | Wilcoxon test | 5.1601E+09 | $n_{Native}$=122678, $n_{Main}$=122678 | $2.2e^{-16}$ |
| 0-fold SNPs | Native vs. Catalonia | $\Theta_w$ | Wilcoxon test | 1.1576E+10 | $n_{Native}$=122678, $n_{Catalonia}$=122678 | $2.2e^{-16}$ |
| 0-fold SNPs | Native vs. Chile | $\Theta_w$ | Wilcoxon test | 8.7194E+09 | $n_{Native}$=122678, $n_{Chile}$=122678 | $2.2e^{-16}$ |
| 0-fold SNPs | Main vs. Catalonia | $\Theta_w$ | Wilcoxon test | 2.8213E+09 | $n_{Main}$=122678, $n_{Catalonia}$=122678 | $2.2e^{-16}$ |
| 0-fold SNPs | Main vs. Chile | $\Theta_w$ | Wilcoxon test | 4.1447E+09 | $n_{Main}$=122678, $n_{Chile}$=122678 | $2.2e^{-16}$ |
| 0-fold | Catalonia vs. | $\Theta_w$ | Wilcoxon | 1.1221E+10 | $n_{Catalonia}$=122678, | $2.2e^{-16}$ |

| | | | | | | |
|---|---|---|---|---|---|---|
| SNPs | Chile | | test | | $n_{Chile}$=122678 | |
| 0-fold SNPs | Native π vs. Native $\Theta_w$ | π & $\Theta_w$ | Wilcoxon test | 1.0239E+09 | $n_{Native(\pi)}$=122678, $n_{Native(\Theta w)}$=122678 | $<2.2e^{-16}$ |
| 0-fold SNPs | Main π vs. Main $\Theta_w$ | π & $\Theta_w$ | Wilcoxon test | 5.2375E+08 | $n_{Main(\pi)}$=122678, $n_{Main(\Theta w)}$=122678 | $<2.2e^{-16}$ |
| 0-fold SNPs | Catalonia π vs. Catalonia $\Theta_w$ | π & $\Theta_w$ | Wilcoxon test | 5.5741E+08 | $n_{Catalonia(\pi)}$=122678, $n_{Catalonia(\Theta w)}$=122678 | $<2.2e^{-16}$ |
| 0-fold SNPs | Chile π vs. Chile $\Theta_w$ | π & $\Theta_w$ | Wilcoxon test | 5.9731E+08 | $n_{Chile(\pi)}$=122678, $n_{Chile(\Theta w)}$=122678 | $<2.2e^{-16}$ |
| 4-fold & 0-fold SNPs | Native 4-fold vs. Native 0-fold | π | Wilcoxon test | 2.7880E+09 | $n_{Native(4fold)}$=59575 $n_{Native(0fold)}$=122678 | $<2.2e^{-16}$ |
| 4-fold & 0-fold SNPs | Main 4-fold vs. Main 0-fold | π | Wilcoxon test | 3.5188E+09 | $n_{Main(4fold)}$=59575 $n_{Main(0fold)}$=122678 | $<2.2e^{-16}$ |
| 4-fold & 0-fold SNPs | Catalonia 4-fold vs. Catalonia 0-fold | π | Wilcoxon test | 3.5883E+09 | $n_{Catalonia(4fold)}$=59575 $n_{Catalonia(0fold)}$=122678 | $<2.2e^{-16}$ |
| 4-fold & 0-fold SNPs | Chile 4-fold vs. Chile 0-fold | π | Wilcoxon test | 3.3955E+09 | $n_{Chile(4fold)}$=59575 $n_{Chile(0fold)}$=122678 | $<2.2e^{-16}$ |
| 4-fold & 0-fold SNPs | Native 4-fold vs. Native 0-fold | $\Theta_w$ | Wilcoxon test | 3.4385E+09 | $n_{Native(4fold)}$=59575 $n_{Native(0fold)}$=122678 | $<2.2e^{-16}$ |
| 4-fold & 0-fold SNPs | Main 4-fold vs. Main 0-fold | $\Theta_w$ | Wilcoxon test | 3.4640E+09 | $n_{Main(4fold)}$=59575 $n_{Main(0fold)}$=122678 | $<2.2e^{-16}$ |
| 4-fold & 0-fold SNPs | Catalonia 4-fold vs. Catalonia 0-fold | $\Theta_w$ | Wilcoxon test | 3.5712E+09 | $n_{Catalonia(4fold)}$=59575 $n_{Catalonia(0fold)}$=122678 | 2.983e-15 |
| 4-fold & 0-fold SNPs | Chile 4-fold vs. Chile 0-fold | $\Theta_w$ | Wilcoxon test | 3.4187E+09 | $n_{Chile(4fold)}$=59575 $n_{Chile(0fold)}$=122678 | $<2.2e^{-16}$ |

**Table S4.** Microsatellite marker diversity expressed as expected heterozygosity ($H_{exp}$) and allelic richness (k'), both on average per five loci. k' is adjusted to a minimum sample size of ten individuals.

| Sample | $H_{exp}$ | $SE_{Hexp}$ | k' | $SE_{k'}$ | N |
|---|---|---|---|---|---|
| Native | 0.9036 | 0.0040 | 9.980 | 0.185 | 10 |
| European Main | 0.4644 | 0.0078 | 3.254 | 0.043 | 10 |
| Catalonia | 0.5710 | 0.0073 | 4.040 | 0.057 | 10 |
| Chile | 0.5426 | 0.0053 | 3.440 | 0.056 | 10 |

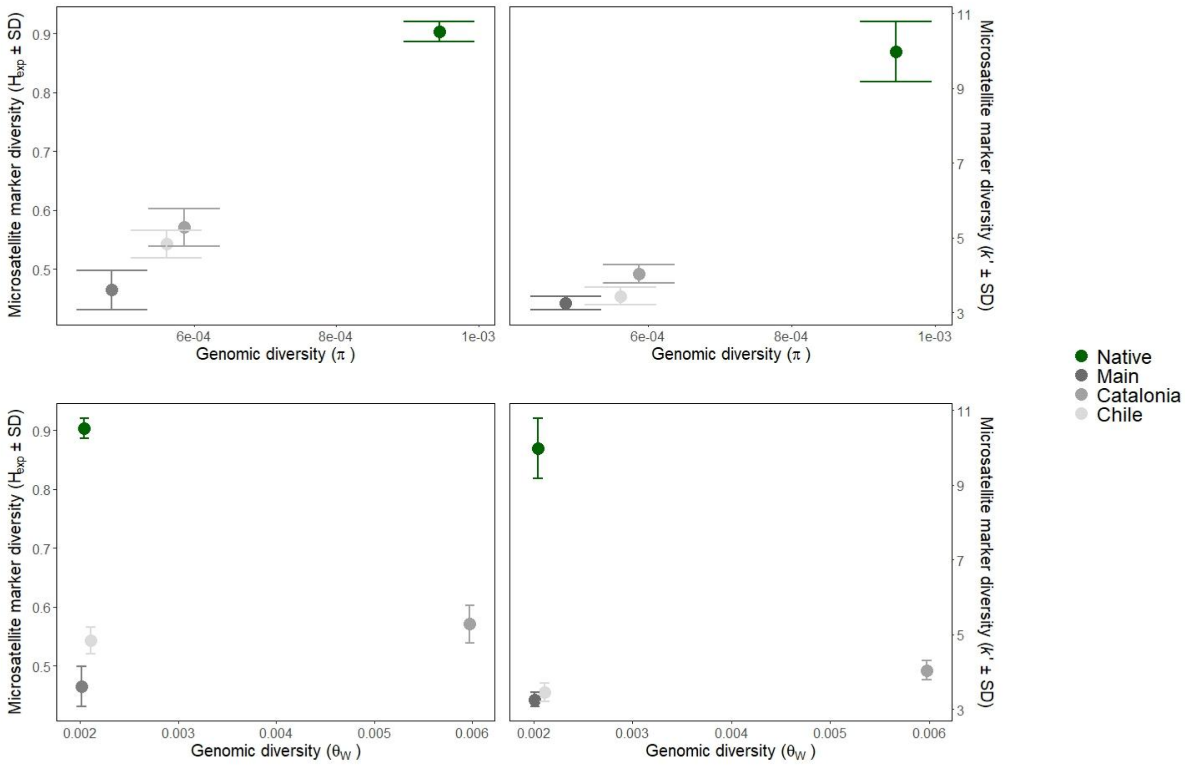


***Figure S5.*** *Genomic diversity (genome-wide average of π and $Θ_w$) comparison with microsatellite marker diversity (average $H_{exp}$ and k' per five loci).*

**Table S6.** Microsatellite marker differentiation as pairwise $F_{ST}$, average per five loci.

| **Sample pair** | **$F_{ST}$** | **SE** | **N** |
|---|---|---|---|
| Native – European Main | 0.2419 | 0.0066 | 10 |
| Native – Catalonia | 0.1945 | 0.0061 | 10 |
| Native – Chile | 0.1902 | 0.0083 | 10 |
| European Main – Catalonia | 0.4565 | 0.0066 | 10 |
| European Main – Chile | 0.3706 | 0.0055 | 10 |
| Chile – Catalonia | 0.3942 | 0.0041 | 10 |

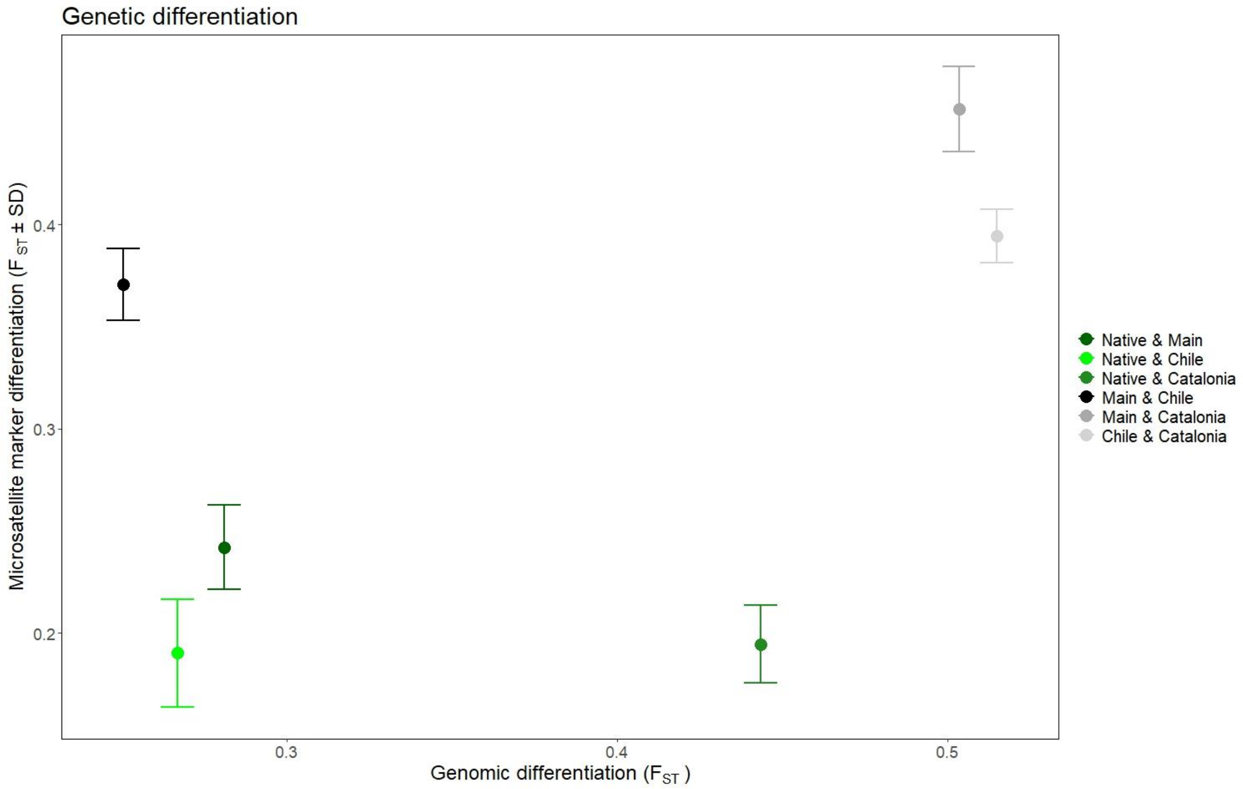


***Figure S7.*** *Genomic differentiation (whole-genome average $F_{ST}$) comparison with microsatellite marker differentiation (average $F_{ST}$ per five loci).*

**Table S8.** Candidate genes for positive selection.

| Gene | Gene ID | Gene | Gene ID | Gene | Gene ID |
|---|---|---|---|---|---|
| LOC105678722 | 105678722 | LOC105671654 | 105671654 | LOC105674409 | 105674409 |
| mRpL54 | 105675240 | LOC105671805 | 105671805 | LOC105674435 | 105674435 |
| LOC105675641 | 105675641 | LOC105671809 | 105671809 | LOC105674437 | 105674437 |
| GluRIB | 105678066 | LOC105671833 | 105671833 | LOC105674476 | 105674476 |
| Scgdelta | 105674460 | LOC105671901 | 105671901 | LOC105674537 | 105674537 |
| Fas1 | 105674287 | LOC105671962 | 105671962 | LOC105674578 | 105674578 |
| lilli | 105669543 | LOC105672037 | 105672037 | LOC105674628 | 105674628 |
| LOC105671159 | 105671159 | LOC105672065 | 105672065 | LOC105674642 | 105674642 |
| LOC105672143 | 105672143 | LOC105672100 | 105672100 | LOC105674782 | 105674782 |
| LOC105668540 | 105668540 | LOC105672102 | 105672102 | LOC105674811 | 105674811 |
| LOC105670711 | 105670711 | LOC105672112 | 105672112 | LOC105674840 | 105674840 |
| SoxN | 105670965 | LOC105672138 | 105672138 | LOC105674880 | 105674880 |
| sowah | 105670960 | LOC105672163 | 105672163 | LOC105674914 | 105674914 |
| LOC105672731 | 105672731 | LOC105672241 | 105672241 | LOC105674982 | 105674982 |
| LOC105669778 | 105669778 | LOC105672268 | 105672268 | LOC105675059 | 105675059 |

| | | | | | |
|---|---|---|---|---|---|
| Ddr | 105678079 | LOC105672553 | 105672553 | LOC105675126 | 105675126 |
| LOC136999726 | 136999726 | LOC105672861 | 105672861 | LOC105675132 | 105675132 |
| LOC105680066 | 105680066 | LOC105672867 | 105672867 | LOC105675161 | 105675161 |
| LOC105672005 | 105672005 | LOC105672889 | 105672889 | LOC105675162 | 105675162 |
| LOC136997244 | 136997244 | LOC105672899 | 105672899 | LOC105675169 | 105675169 |
| Prosap | 105672750 | LOC105673042 | 105673042 | LOC105675175 | 105675175 |
| LOC105670217 | 105670217 | LOC105673158 | 105673158 | LOC105675189 | 105675189 |
| tn | 105674874 | LOC105673241 | 105673241 | LOC105675222 | 105675222 |
| LOC105675255 | 105675255 | LOC105673414 | 105673414 | LOC105675239 | 105675239 |
| LOC136997196 | 136997196 | LOC105673430 | 105673430 | LOC105675249 | 105675249 |
| Shrm | 105676893 | LOC105673480 | 105673480 | LOC105675316 | 105675316 |
| LOC105668956 | 105668956 | LOC105673485 | 105673485 | LOC105675357 | 105675357 |
| LOC105670963 | 105670963 | LOC105673872 | 105673872 | LOC105675365 | 105675365 |
| LOC105671998 | 105671998 | LOC105674241 | 105674241 | LOC105675445 | 105675445 |
| LOC105672277 | 105672277 | LOC105674349 | 105674349 | LOC105675446 | 105675446 |
| LOC105679894 | 105679894 | LOC105674432 | 105674432 | LOC105675525 | 105675525 |
| LOC136998128 | 136998128 | LOC105674536 | 105674536 | LOC105675593 | 105675593 |
| dtn | 105678176 | LOC105674596 | 105674596 | LOC105675642 | 105675642 |
| LOC105669009 | 105669009 | LOC105674630 | 105674630 | LOC105675694 | 105675694 |
| LOC105677977 | 105677977 | LOC105674684 | 105674684 | LOC105675830 | 105675830 |
| LOC105669582 | 105669582 | LOC105674817 | 105674817 | LOC105675837 | 105675837 |
| LOC105675252 | 105675252 | LOC105674860 | 105674860 | LOC105675875 | 105675875 |
| LOC105676847 | 105676847 | LOC105675098 | 105675098 | LOC105675915 | 105675915 |
| LOC136997100 | 136997100 | LOC105675136 | 105675136 | LOC105675959 | 105675959 |
| LOC136999427 | 136999427 | LOC105675234 | 105675234 | LOC105676069 | 105676069 |
| LOC105672068 | 105672068 | LOC105675236 | 105675236 | LOC105676070 | 105676070 |
| Lim1 | 105679908 | LOC105675373 | 105675373 | LOC105676102 | 105676102 |
| LOC105672015 | 105672015 | LOC105675427 | 105675427 | LOC105676118 | 105676118 |
| stet | 105676423 | LOC105675468 | 105675468 | LOC105676274 | 105676274 |
| Fs | 105675232 | LOC105675816 | 105675816 | LOC105676365 | 105676365 |
| LOC105668537 | 105668537 | LOC105675855 | 105675855 | LOC105676378 | 105676378 |
| LOC105670446 | 105670446 | LOC105676137 | 105676137 | LOC105676404 | 105676404 |
| LOC105679896 | 105679896 | LOC105676184 | 105676184 | LOC105676468 | 105676468 |
| Mf | 105667323 | LOC105676399 | 105676399 | LOC105676470 | 105676470 |
| ds | 105672238 | LOC105676432 | 105676432 | LOC105676479 | 105676479 |
| LOC105672197 | 105672197 | LOC105676443 | 105676443 | LOC105676550 | 105676550 |
| LOC105674883 | 105674883 | LOC105676454 | 105676454 | LOC105676593 | 105676593 |
| bru3 | 105671757 | LOC105676570 | 105676570 | LOC105676633 | 105676633 |
| Cad74A | 105675653 | LOC105676620 | 105676620 | LOC105676645 | 105676645 |
| GckIII | 105667667 | LOC105676639 | 105676639 | LOC105676651 | 105676651 |
| LOC105670160 | 105670160 | LOC105676664 | 105676664 | LOC105676653 | 105676653 |

| | | | | | |
|---|---|---|---|---|---|
| LOC105672946 | 105672946 | LOC105676668 | 105676668 | LOC105676654 | 105676654 |
| LOC105674436 | 105674436 | LOC105677094 | 105677094 | LOC105676657 | 105676657 |
| LOC105675250 | 105675250 | LOC105677108 | 105677108 | LOC105676658 | 105676658 |
| LOC105676396 | 105676396 | LOC105677137 | 105677137 | LOC105676659 | 105676659 |
| LOC105679779 | 105679779 | LOC105677139 | 105677139 | LOC105676663 | 105676663 |
| LOC137000536 | 137000536 | LOC105677157 | 105677157 | LOC105676688 | 105676688 |
| LOC137000540 | 137000540 | LOC105677193 | 105677193 | LOC105676735 | 105676735 |
| Syn1 | 105674884 | LOC105677342 | 105677342 | LOC105676739 | 105676739 |
| dlp | 105677985 | LOC105677589 | 105677589 | LOC105676742 | 105676742 |
| vn | 105668495 | LOC105677812 | 105677812 | LOC105676960 | 105676960 |
| HisT | 105671984 | LOC105677858 | 105677858 | LOC105677055 | 105677055 |
| LOC105667666 | 105667666 | LOC105678290 | 105678290 | LOC105677116 | 105677116 |
| LOC105667782 | 105667782 | LOC105678319 | 105678319 | LOC105677124 | 105677124 |
| LOC105673538 | 105673538 | LOC105678455 | 105678455 | LOC105677148 | 105677148 |
| LOC105675049 | 105675049 | LOC105678465 | 105678465 | LOC105677195 | 105677195 |
| LOC136999630 | 136999630 | LOC105678466 | 105678466 | LOC105677212 | 105677212 |
| LOC137000479 | 137000479 | LOC105678601 | 105678601 | LOC105677310 | 105677310 |
| alpha-Man-IIb | 105674429 | LOC105678667 | 105678667 | LOC105677318 | 105677318 |
| inaE | 105668919 | LOC105678706 | 105678706 | LOC105677344 | 105677344 |
| kug | 105668354 | LOC105678707 | 105678707 | LOC105677432 | 105677432 |
| sls | 105670419 | LOC105678718 | 105678718 | LOC105677448 | 105677448 |
| sns | 105674585 | LOC105678761 | 105678761 | LOC105677508 | 105677508 |
| Ae2 | 105673500 | LOC105678799 | 105678799 | LOC105677579 | 105677579 |
| Apoltp | 105679739 | LOC105678865 | 105678865 | LOC105677953 | 105677953 |
| LOC105667781 | 105667781 | LOC105679055 | 105679055 | LOC105677984 | 105677984 |
| LOC105668274 | 105668274 | LOC105679131 | 105679131 | LOC105677989 | 105677989 |
| LOC105668961 | 105668961 | LOC105679504 | 105679504 | LOC105678038 | 105678038 |
| LOC105669058 | 105669058 | LOC105679773 | 105679773 | LOC105678041 | 105678041 |
| LOC105669447 | 105669447 | LOC105679782 | 105679782 | LOC105678070 | 105678070 |
| LOC105670961 | 105670961 | LOC105679837 | 105679837 | LOC105678081 | 105678081 |
| LOC105678633 | 105678633 | LOC105679969 | 105679969 | LOC105678110 | 105678110 |
| LOC136998114 | 136998114 | LOC105679993 | 105679993 | LOC105678130 | 105678130 |
| milt | 105679946 | LOC105680017 | 105680017 | LOC105678131 | 105678131 |
| wb | 105668754 | LOC136996908 | 136996908 | LOC105678152 | 105678152 |
| LOC105668862 | 105668862 | LOC136996946 | 136996946 | LOC105678174 | 105678174 |
| LOC105668880 | 105668880 | LOC136997036 | 136997036 | LOC105678177 | 105678177 |
| LOC105668952 | 105668952 | LOC136997076 | 136997076 | LOC105678197 | 105678197 |
| LOC105669593 | 105669593 | LOC136997151 | 136997151 | LOC105678212 | 105678212 |
| LOC105670148 | 105670148 | LOC136997190 | 136997190 | LOC105678230 | 105678230 |
| LOC105673149 | 105673149 | LOC136997315 | 136997315 | LOC105678312 | 105678312 |
| LOC105673369 | 105673369 | LOC136997398 | 136997398 | LOC105678327 | 105678327 |

| | | | | | |
|---|---|---|---|---|---|
| LOC105674360 | 105674360 | LOC136997746 | 136997746 | LOC105678348 | 105678348 |
| LOC105675100 | 105675100 | LOC136997804 | 136997804 | LOC105678360 | 105678360 |
| LOC105676698 | 105676698 | LOC136997950 | 136997950 | LOC105678361 | 105678361 |
| LOC105677337 | 105677337 | LOC136998049 | 136998049 | LOC105678362 | 105678362 |
| LOC137000775 | 137000775 | LOC136999265 | 136999265 | LOC105678409 | 105678409 |
| LanA | 105672030 | LOC136999343 | 136999343 | LOC105678422 | 105678422 |
| Liprin-gamma | 105672672 | LOC136999548 | 136999548 | LOC105678452 | 105678452 |
| Rbp6 | 105670561 | LOC136999956 | 136999956 | LOC105678453 | 105678453 |
| Sp1 | 105672900 | LOC136999988 | 136999988 | LOC105678480 | 105678480 |
| Trpgamma | 105669094 | LOC137000135 | 137000135 | LOC105678619 | 105678619 |
| atk | 105670977 | LOC137000740 | 137000740 | LOC105678626 | 105678626 |
| dnc | 105678731 | LOC137000774 | 137000774 | LOC105678704 | 105678704 |
| exex | 105677110 | LOC137001045 | 137001045 | LOC105678713 | 105678713 |
| if | 105670985 | LOC137001108 | 137001108 | LOC105678717 | 105678717 |
| mirr | 105670959 | LOC137001235 | 137001235 | LOC105678741 | 105678741 |
| Hr38 | 105676504 | LOC137001433 | 137001433 | LOC105678809 | 105678809 |
| LOC105667445 | 105667445 | LOC137001681 | 137001681 | LOC105678847 | 105678847 |
| LOC105667909 | 105667909 | LOC137001906 | 137001906 | LOC105678888 | 105678888 |
| LOC105668392 | 105668392 | Lis-1 | 105668399 | LOC105678899 | 105678899 |
| LOC105668524 | 105668524 | Msp300 | 105675233 | LOC105678906 | 105678906 |
| LOC105668533 | 105668533 | Nmdar2 | 105670141 | LOC105678911 | 105678911 |
| LOC105671659 | 105671659 | Nrx-1 | 105678852 | LOC105678969 | 105678969 |
| LOC105671821 | 105671821 | Ntan1 | 105674442 | LOC105678973 | 105678973 |
| LOC105671981 | 105671981 | P32 | 105677171 | LOC105679022 | 105679022 |
| LOC105672009 | 105672009 | Parp1 | 105676389 | LOC105679042 | 105679042 |
| LOC105675238 | 105675238 | Pax | 105677158 | LOC105679062 | 105679062 |
| LOC105676114 | 105676114 | Pgant9 | 105670712 | LOC105679084 | 105679084 |
| LOC105677968 | 105677968 | Pka-R1 | 105668802 | LOC105679095 | 105679095 |
| LOC105678008 | 105678008 | PlexA | 105678134 | LOC105679134 | 105679134 |
| LOC105678034 | 105678034 | PolE1 | 105677922 | LOC105679181 | 105679181 |
| LOC105678057 | 105678057 | Ptp69D | 105668417 | LOC105679337 | 105679337 |
| LOC105679013 | 105679013 | Ptp99A | 105669580 | LOC105679569 | 105679569 |
| LOC105679130 | 105679130 | Qsox1 | 105675318 | LOC105679596 | 105679596 |
| LOC137000684 | 137000684 | Reck | 105671334 | LOC105679623 | 105679623 |
| RhoGAP15B | 105678602 | RhoGEF3 | 105677499 | LOC105679646 | 105679646 |
| cindr | 105669756 | Rim2 | 105678137 | LOC105679687 | 105679687 |
| fmt | 105672049 | SK | 105673813 | LOC105679726 | 105679726 |
| mav | 105669559 | Sema5c | 105672199 | LOC105679765 | 105679765 |
| pea | 105670279 | Snap25 | 105677192 | LOC105679818 | 105679818 |
| pnut | 105677197 | Toll-6 | 105674053 | LOC105679824 | 105679824 |
| sd | 105676108 | Trax | 105672046 | LOC105679835 | 105679835 |

| | | | | | |
|---|---|---|---|---|---|
| v | 105680137 | Trim9 | 105671481 | LOC105679840 | 105679840 |
| zip | 105676387 | TrissinR | 105676794 | LOC105679858 | 105679858 |
| Cals | 105679447 | Trl | 105680065 | LOC105679958 | 105679958 |
| Fas3 | 105671823 | Trpm | 105679369 | LOC105679975 | 105679975 |
| GluClalpha | 105671992 | Unc-76 | 105675769 | LOC105680005 | 105680005 |
| Klp31E | 105667326 | Utx | 105672366 | LOC105680014 | 105680014 |
| LOC105667438 | 105667438 | Wdr81 | 105671370 | LOC105680016 | 105680016 |
| LOC105667821 | 105667821 | acj6 | 105672811 | LOC105680018 | 105680018 |
| LOC105669746 | 105669746 | bif | 105673269 | LOC105680020 | 105680020 |
| LOC105669910 | 105669910 | bsk | 105670179 | LOC105680086 | 105680086 |
| LOC105670149 | 105670149 | cher | 105667827 | LOC105680164 | 105680164 |
| LOC105671698 | 105671698 | crb | 105670455 | LOC136996894 | 136996894 |
| LOC105671730 | 105671730 | dlg1 | 105677469 | LOC136996923 | 136996923 |
| LOC105672220 | 105672220 | dmpd | 105668212 | LOC136996954 | 136996954 |
| LOC105672709 | 105672709 | dpr12 | 105674107 | LOC136997057 | 136997057 |
| LOC105674045 | 105674045 | eEF2 | 105678604 | LOC136997065 | 136997065 |
| LOC105674138 | 105674138 | ena | 105670398 | LOC136997101 | 136997101 |
| LOC105674446 | 105674446 | fry | 105675416 | LOC136997180 | 136997180 |
| LOC105674609 | 105674609 | fs(1)Ya | 105679906 | LOC136997229 | 136997229 |
| LOC105674723 | 105674723 | fwd | 105677540 | LOC136997245 | 136997245 |
| LOC105674936 | 105674936 | heph | 105678414 | LOC136997288 | 136997288 |
| LOC105675912 | 105675912 | hop | 105677920 | LOC136997299 | 136997299 |
| LOC105675932 | 105675932 | ich | 105677349 | LOC136997325 | 136997325 |
| LOC105676622 | 105676622 | imd | 105672003 | LOC136997361 | 136997361 |
| LOC105676646 | 105676646 | kmr | 105674224 | LOC136997375 | 136997375 |
| LOC105676673 | 105676673 | kuz | 105675974 | LOC136997397 | 136997397 |
| LOC105676845 | 105676845 | lola | 105674339 | LOC136997407 | 136997407 |
| LOC105678238 | 105678238 | mTerf5 | 105667736 | LOC136997527 | 136997527 |
| LOC105678650 | 105678650 | mib1 | 105677910 | LOC136997541 | 136997541 |
| LOC105679791 | 105679791 | mtd | 105669392 | LOC136997631 | 136997631 |
| LOC105679882 | 105679882 | nej | 105669562 | LOC136997675 | 136997675 |
| LOC136998039 | 136998039 | pxb | 105670734 | LOC136997744 | 136997744 |
| LOC136999401 | 136999401 | pygo | 105672898 | LOC136997772 | 136997772 |
| LOC137000466 | 137000466 | raskol | 105668128 | LOC136997780 | 136997780 |
| LOC137000480 | 137000480 | rdx | 105679641 | LOC136997781 | 136997781 |
| Mdr49 | 105680135 | rg | 105668297 | LOC136997816 | 136997816 |
| Pdk1 | 105672857 | sano | 105672914 | LOC136997843 | 136997843 |
| Smr | 105673525 | sdk | 105678005 | LOC136997966 | 136997966 |
| beta'COP | 105674430 | sima | 105677168 | LOC136997992 | 136997992 |
| cdi | 105671834 | spri | 105667764 | LOC136997998 | 136997998 |
| dally | 105671660 | sv | 105675806 | LOC136998009 | 136998009 |

| | | | | | |
|---|---|---|---|---|---|
| gish | 105679530 | svp | 105670852 | LOC136998044 | 136998044 |
| p130CAS | 105676498 | tweek | 105671293 | LOC136998120 | 136998120 |
| sha | 105677990 | zda | 105671461 | LOC136998133 | 136998133 |
| tei | 105674123 | 5-HT2B | 105667979 | LOC136998413 | 136998413 |
| ths | 105672274 | AIF | 105678244 | LOC136998472 | 136998472 |
| wdp | 105667683 | Acf | 105667873 | LOC136998519 | 136998519 |
| Blimp-1 | 105668430 | Adsl | 105673778 | LOC136998561 | 136998561 |
| CadN | 105674346 | Adss | 105676507 | LOC136998643 | 136998643 |
| DIP2 | 105673477 | Ald1 | 105673579 | LOC136998670 | 136998670 |
| Hers | 105676362 | Amph | 105678981 | LOC136998835 | 136998835 |
| Itpr | 105670881 | Archease | 105670449 | LOC136998856 | 136998856 |
| LOC105667349 | 105667349 | ArfGAP3 | 105678330 | LOC136998869 | 136998869 |
| LOC105668432 | 105668432 | Arfip | 105667596 | LOC136999259 | 136999259 |
| LOC105668497 | 105668497 | Asap | 105676503 | LOC136999264 | 136999264 |
| LOC105669532 | 105669532 | Awh | 105670079 | LOC136999314 | 136999314 |
| LOC105669591 | 105669591 | Bap170 | 105675939 | LOC136999334 | 136999334 |
| LOC105669665 | 105669665 | Brf | 105673584 | LOC136999348 | 136999348 |
| LOC105669681 | 105669681 | CTCF | 105675224 | LOC136999481 | 136999481 |
| LOC105670081 | 105670081 | Camta | 105674935 | LOC136999489 | 136999489 |
| LOC105670224 | 105670224 | Cat | 105676169 | LOC136999684 | 136999684 |
| LOC105670479 | 105670479 | Ccdc114 | 105670170 | LOC136999697 | 136999697 |
| LOC105670483 | 105670483 | CdGAPr | 105678433 | LOC136999785 | 136999785 |
| LOC105670491 | 105670491 | CrebA | 105675033 | LOC136999990 | 136999990 |
| LOC105671259 | 105671259 | CycB3 | 105673764 | LOC136999992 | 136999992 |
| LOC105671360 | 105671360 | CycG | 105668922 | LOC137000030 | 137000030 |
| LOC105672019 | 105672019 | CycT | 105669763 | LOC137000044 | 137000044 |
| LOC105672165 | 105672165 | DCTN6-p27 | 105679003 | LOC137000093 | 137000093 |
| LOC105672326 | 105672326 | DUBAI | 105677118 | LOC137000106 | 137000106 |
| LOC105672801 | 105672801 | Dab | 105677489 | LOC137000118 | 137000118 |
| LOC105673142 | 105673142 | Delta | 105671450 | LOC137000152 | 137000152 |
| LOC105673159 | 105673159 | Dh31 | 105679825 | LOC137000197 | 137000197 |
| LOC105673481 | 105673481 | Dhc64C | 105676072 | LOC137000335 | 137000335 |
| LOC105673771 | 105673771 | Dhc98D | 105671582 | LOC137000368 | 137000368 |
| LOC105674282 | 105674282 | Doa | 105668220 | LOC137000384 | 137000384 |
| LOC105674357 | 105674357 | Dpp10 | 105669452 | LOC137000515 | 137000515 |
| LOC105675122 | 105675122 | Drgx | 105671965 | LOC137000522 | 137000522 |
| LOC105675522 | 105675522 | Drgx | 105671965 | LOC137000551 | 137000551 |
| LOC105676656 | 105676656 | Eip75B | 105674872 | LOC137000644 | 137000644 |
| LOC105676823 | 105676823 | EndoA | 105670622 | LOC137000654 | 137000654 |
| LOC105676868 | 105676868 | Ets65A | 105676413 | LOC137000662 | 137000662 |
| LOC105678625 | 105678625 | Ets96B | 105679624 | LOC137000783 | 137000783 |

| | | | | | |
|---|---|---|---|---|---|
| LOC105678710 | 105678710 | Exd2 | 105672596 | LOC137000814 | 137000814 |
| LOC105678744 | 105678744 | Flo2 | 105669524 | LOC137000980 | 137000980 |
| LOC105679089 | 105679089 | Galk | 105677111 | LOC137001010 | 137001010 |
| LOC105679456 | 105679456 | Galphao | 105678675 | LOC137001143 | 137001143 |
| LOC105679582 | 105679582 | Gfrl | 105679160 | LOC137001163 | 137001163 |
| LOC105679614 | 105679614 | Gga | 105675582 | LOC137001280 | 137001280 |
| LOC105679685 | 105679685 | Hen1 | 105679398 | LOC137001417 | 137001417 |
| LOC136996988 | 136996988 | Hira | 105675235 | LOC137001904 | 137001904 |
| LOC136997099 | 136997099 | Hml | 105670297 | LOC137001926 | 137001926 |
| LOC136997526 | 136997526 | ITP | 105671648 | LOC137001941 | 137001941 |
| LOC136997739 | 136997739 | Iml1 | 105675544 | Lasp | 105675091 |
| LOC136997763 | 136997763 | IntS14 | 105675088 | Lcch3 | 105667704 |
| LOC136999253 | 136999253 | Isha | 105675004 | Lgr3 | 105667860 |
| LOC137000986 | 137000986 | Jhbp1 | 105670444 | Lim3 | 105674489 |
| Lrch | 105674127 | KCNQ | 105674186 | Lon | 105673088 |
| Patronin | 105668761 | Kdm2 | 105674731 | Lst8 | 105675583 |
| Task6 | 105674019 | Kdm3 | 105671081 | M6 | 105679891 |
| Ten-a | 105668275 | LOC105667289 | 105667289 | MED20 | 105673405 |
| Tis11 | 105669901 | LOC105667319 | 105667319 | MED23 | 105674457 |
| Vps13D | 105670276 | LOC105667419 | 105667419 | MESR6 | 105673939 |
| aPKC | 105669556 | LOC105667432 | 105667432 | MFS16 | 105668931 |
| chb | 105671218 | LOC105667443 | 105667443 | MSBP | 105669576 |
| dachs | 105674808 | LOC105667446 | 105667446 | Mat89Ba | 105667786 |
| heca | 105675434 | LOC105667449 | 105667449 | Mgat4a | 105672071 |
| hiw | 105675379 | LOC105667559 | 105667559 | Mlp84B | 105678133 |
| jing | 105672755 | LOC105667578 | 105667578 | Mob2 | 105677000 |
| mthl1 | 105678795 | LOC105667583 | 105667583 | Mppe | 105674300 |
| nrm | 105669214 | LOC105667629 | 105667629 | Myb | 105668525 |
| peb | 105677699 | LOC105667632 | 105667632 | Myo31DF | 105674535 |
| qin | 105668992 | LOC105667644 | 105667644 | Myo61F | 105679915 |
| sli | 105670472 | LOC105667647 | 105667647 | NKCC | 105675372 |
| sxc | 105674563 | LOC105667650 | 105667650 | Nca | 105677509 |
| Best2 | 105676966 | LOC105667655 | 105667655 | Ncc69 | 105669100 |
| Cyp18a1 | 105675524 | LOC105667702 | 105667702 | Notum | 105678106 |
| Cyp303a1 | 105672072 | LOC105667703 | 105667703 | Np | 105674926 |
| Dlg5 | 105671699 | LOC105667758 | 105667758 | Ns1 | 105671784 |
| Dscam3 | 105671699 | LOC105667787 | 105667787 | Nup153 | 105678042 |
| Glut4EF | 105678792 | LOC105667851 | 105667851 | Octalpha2R | 105675384 |
| LOC105667440 | 105667440 | LOC105667853 | 105667853 | Orc3 | 105674288 |
| LOC105667442 | 105667442 | LOC105667866 | 105667866 | Ork1 | 105673406 |
| LOC105667715 | 105667715 | LOC105667876 | 105667876 | Oseg6 | 105669573 |

| | | | | | |
|---|---|---|---|---|---|
| LOC105668652 | 105668652 | LOC105667912 | 105667912 | PIG-S | 105673656 |
| LOC105668939 | 105668939 | LOC105667915 | 105667915 | PRAS40 | 105673603 |
| LOC105668964 | 105668964 | LOC105668027 | 105668027 | Pa1 | 105679667 |
| LOC105668997 | 105668997 | LOC105668215 | 105668215 | Pdp | 105673552 |
| LOC105669315 | 105669315 | LOC105668247 | 105668247 | Pdp1 | 105669686 |
| LOC105669417 | 105669417 | LOC105668260 | 105668260 | Pgant2 | 105670106 |
| LOC105669596 | 105669596 | LOC105668264 | 105668264 | Pgi | 105669444 |
| LOC105669909 | 105669909 | LOC105668326 | 105668326 | Phb1 | 105670973 |
| LOC105670496 | 105670496 | LOC105668373 | 105668373 | Pino | 105675713 |
| LOC105670558 | 105670558 | LOC105668556 | 105668556 | Pp2C1 | 105667663 |
| LOC105670597 | 105670597 | LOC105668583 | 105668583 | Ptr | 105675520 |
| LOC105670614 | 105670614 | LOC105668599 | 105668599 | REPTOR-BP | 105667348 |
| LOC105671270 | 105671270 | LOC105668682 | 105668682 | RPA1 | 105677186 |
| LOC105671375 | 105671375 | LOC105668686 | 105668686 | Rab35 | 105673712 |
| LOC105671377 | 105671377 | LOC105668719 | 105668719 | RasGAP1 | 105675697 |
| LOC105671581 | 105671581 | LOC105668725 | 105668725 | Rbfox1 | 105673888 |
| LOC105672035 | 105672035 | LOC105668796 | 105668796 | Rdl | 105678045 |
| LOC105672043 | 105672043 | LOC105668842 | 105668842 | Remo | 105671714 |
| LOC105672530 | 105672530 | LOC105668866 | 105668866 | Rgk2 | 105668439 |
| LOC105672694 | 105672694 | LOC105668905 | 105668905 | Rint1 | 105671371 |
| LOC105673842 | 105673842 | LOC105668925 | 105668925 | RpL11 | 105679959 |
| LOC105674041 | 105674041 | LOC105668932 | 105668932 | RpS19a | 105676594 |
| LOC105674058 | 105674058 | LOC105668950 | 105668950 | Rtnl1 | 105669458 |
| LOC105674444 | 105674444 | LOC105669027 | 105669027 | RyR | 105669060 |
| LOC105675102 | 105675102 | LOC105669120 | 105669120 | SIDL | 105674117 |
| LOC105675117 | 105675117 | LOC105669150 | 105669150 | SamDC | 105676778 |
| LOC105675158 | 105675158 | LOC105669159 | 105669159 | Sara | 105671321 |
| LOC105675173 | 105675173 | LOC105669433 | 105669433 | Sbf | 105667711 |
| LOC105675241 | 105675241 | LOC105669449 | 105669449 | Sdc | 105679794 |
| LOC105675256 | 105675256 | LOC105669455 | 105669455 | Sema2a | 105670029 |
| LOC105676139 | 105676139 | LOC105669469 | 105669469 | Sfxn2 | 105675740 |
| LOC105676519 | 105676519 | LOC105669497 | 105669497 | Sgf11 | 105677188 |
| LOC105676717 | 105676717 | LOC105669520 | 105669520 | Sgf29 | 105670910 |
| LOC105676884 | 105676884 | LOC105669551 | 105669551 | Shal | 105667828 |
| LOC105677127 | 105677127 | LOC105669683 | 105669683 | Smu1 | 105671715 |
| LOC105678048 | 105678048 | LOC105669695 | 105669695 | Smyd4-2 | 105678743 |
| LOC105678143 | 105678143 | LOC105669740 | 105669740 | Syn2 | 105668854 |
| LOC105678752 | 105678752 | LOC105669742 | 105669742 | Syx18 | 105672361 |
| LOC105679047 | 105679047 | LOC105669745 | 105669745 | Syx1A | 105673781 |
| LOC105679106 | 105679106 | LOC105669782 | 105669782 | TSG101 | 105672050 |
| LOC105679373 | 105679373 | LOC105669877 | 105669877 | Tctp | 105677972 |

| | | | | | |
|---|---|---|---|---|---|
| LOC105679811 | 105679811 | LOC105669882 | 105669882 | Teh1 | 105667748 |
| LOC105680036 | 105680036 | LOC105669911 | 105669911 | Ten-m | 105669999 |
| LOC136996865 | 136996865 | LOC105669964 | 105669964 | Top3beta | 105667342 |
| LOC136997098 | 136997098 | LOC105669986 | 105669986 | Tpst | 105673267 |
| LOC136997203 | 136997203 | LOC105670017 | 105670017 | Trmt61 | 105673617 |
| LOC136997628 | 136997628 | LOC105670031 | 105670031 | Tsp42Ed | 105669779 |
| LOC136998453 | 136998453 | LOC105670036 | 105670036 | U2af50 | 105669129 |
| LOC136998542 | 136998542 | LOC105670037 | 105670037 | UBL3 | 105673093 |
| LOC136999553 | 136999553 | LOC105670042 | 105670042 | Ube4A | 105673568 |
| LOC137000428 | 137000428 | LOC105670051 | 105670051 | Upf3 | 105677512 |
| LOC137000467 | 137000467 | LOC105670058 | 105670058 | Vps25 | 105678620 |
| LOC137000523 | 137000523 | LOC105670059 | 105670059 | Vps26 | 105676377 |
| LOC137000776 | 137000776 | LOC105670080 | 105670080 | ZnT63C | 105671732 |
| LOC137001058 | 137001058 | LOC105670102 | 105670102 | alpha-Spec | 105670534 |
| Lar | 105674730 | LOC105670161 | 105670161 | aralar1 | 105676467 |
| Lmpt | 105668256 | LOC105670207 | 105670207 | arg | 105671201 |
| Picot | 105673596 | LOC105670281 | 105670281 | ave | 105680182 |
| Pkn | 105670964 | LOC105670299 | 105670299 | bark | 105679489 |
| Polr3E | 105670508 | LOC105670430 | 105670430 | bbx | 105676586 |
| S6KL | 105670065 | LOC105670435 | 105670435 | c12.1 | 105673798 |
| Sf3a2 | 105677609 | LOC105670442 | 105670442 | caly | 105676802 |
| Sh | 105674496 | LOC105670450 | 105670450 | chm | 105677763 |
| Sidpn | 105675019 | LOC105670456 | 105670456 | chn | 105672894 |
| Ssdp | 105668954 | LOC105670460 | 105670460 | dia | 105673157 |
| Tm1 | 105675101 | LOC105670473 | 105670473 | dyl | 105677336 |
| amrt | 105668390 | LOC105670474 | 105670474 | dysf | 105672038 |
| bbg | 105675822 | LOC105670527 | 105670527 | eag | 105675110 |
| brat | 105670830 | LOC105670530 | 105670530 | emp | 105678693 |
| btv | 105670328 | LOC105670535 | 105670535 | enc | 105671418 |
| ct | 105676394 | LOC105670596 | 105670596 | enok | 105676965 |
| disco-r | 105676987 | LOC105670626 | 105670626 | ex | 105673549 |
| dnr1 | 105672154 | LOC105670671 | 105670671 | exd | 105669672 |
| eIF5B | 105669485 | LOC105670836 | 105670836 | fbp | 105671994 |
| east | 105675109 | LOC105670905 | 105670905 | fs(2)ltoPP43 | 105676830 |
| fng | 105675165 | LOC105670972 | 105670972 | fuss | 105670679 |
| fz2 | 105676477 | LOC105670983 | 105670983 | gudu | 105679725 |
| gukh | 105678421 | LOC105670994 | 105670994 | kcc | 105671636 |
| inaF-D | 105672070 | LOC105671170 | 105671170 | kdn | 105668941 |
| mAChR-B | 105677091 | LOC105671181 | 105671181 | kel | 105667590 |
| mlt | 105676100 | LOC105671183 | 105671183 | ken | 105677338 |
| nudC | 105676403 | LOC105671219 | 105671219 | kn | 105672285 |

| | | | | | |
|---|---|---|---|---|---|
| pdm3 | 105670152 | LOC105671609 | 105671609 | kst | 105670477 |
| pnt | 105669422 | LOC105671621 | 105671621 | kto | 105670550 |
| sdt | 105676121 | LOC105671641 | 105671641 | l(2)05287 | 105667752 |
| tmod | 105674290 | LOC105671642 | 105671642 | l(2)dtl | 105670334 |
| tst | 105673731 | LOC105671696 | 105671696 | lace | 105669456 |
| twz | 105678040 | LOC105671792 | 105671792 | ldd | 105680133 |
| Actn | 105673416 | LOC105671858 | 105671858 | lov | 105670832 |
| Arms | 105674792 | LOC105671985 | 105671985 | lqf | 105678182 |
| Atpalpha | 105675982 | LOC105671990 | 105671990 | mIF2 | 105680001 |
| CAP | 105671996 | LOC105672000 | 105672000 | mRpL24 | 105672753 |
| Calx | 105670518 | LOC105672021 | 105672021 | mRpL34 | 105667868 |
| Ccdc56 | 105672677 | LOC105672101 | 105672101 | mRpL52 | 105668251 |
| Cep97 | 105669578 | LOC105672118 | 105672118 | mbo | 105676599 |
| Chd64 | 105674141 | LOC105672137 | 105672137 | mib2 | 105674504 |
| Coq6 | 105677136 | LOC105672141 | 105672141 | mnb | 105667968 |
| Cph | 105676445 | LOC105672153 | 105672153 | msl-3 | 105679608 |
| Dhit | 105672896 | LOC105672208 | 105672208 | mura | 105675317 |
| Dys | 105674116 | LOC105672253 | 105672253 | mys | 105672375 |
| Eph | 105677044 | LOC105672299 | 105672299 | nAChRalpha4 | 105678965 |
| Fife | 105672674 | LOC105672304 | 105672304 | ome | 105674399 |
| Gli | 105674132 | LOC105672327 | 105672327 | osk | 105676478 |
| HDAC4 | 105673647 | LOC105672359 | 105672359 | p | 105673113 |
| HGTX | 105677901 | LOC105672535 | 105672535 | p120ctn | 105676728 |
| HUWE1 | 105676126 | LOC105672573 | 105672573 | pain | 105673141 |
| HnRNP-K | 105677937 | LOC105672597 | 105672597 | phtm | 105675509 |
| KaiR1D | 105678101 | LOC105672712 | 105672712 | pio | 105671497 |
| LOC105667332 | 105667332 | LOC105672732 | 105672732 | poe | 105678791 |
| LOC105667335 | 105667335 | LOC105672826 | 105672826 | pont | 105668903 |
| LOC105667436 | 105667436 | LOC105672858 | 105672858 | ps | 105677151 |
| LOC105667448 | 105667448 | LOC105672892 | 105672892 | qtc | 105680003 |
| LOC105667551 | 105667551 | LOC105672893 | 105672893 | raptor | 105671423 |
| LOC105667894 | 105667894 | LOC105672955 | 105672955 | rdgB | 105677427 |
| LOC105668043 | 105668043 | LOC105673069 | 105673069 | rempA | 105673275 |
| LOC105668273 | 105668273 | LOC105673083 | 105673083 | rho-5 | 105674917 |
| LOC105668304 | 105668304 | LOC105673107 | 105673107 | rho-6 | 105669411 |
| LOC105668370 | 105668370 | LOC105673116 | 105673116 | rsh | 105676558 |
| LOC105668388 | 105668388 | LOC105673179 | 105673179 | shep | 105673189 |
| LOC105668633 | 105668633 | LOC105673218 | 105673218 | sigmar | 105670645 |
| LOC105668687 | 105668687 | LOC105673237 | 105673237 | siz | 105667799 |
| LOC105668861 | 105668861 | LOC105673240 | 105673240 | sm | 105679114 |
| LOC105668907 | 105668907 | LOC105673248 | 105673248 | smash | 105667735 |

| | | | | | |
|---|---|---|---|---|---|
| LOC105668914 | 105668914 | LOC105673310 | 105673310 | snRNP-U1-70K | 105674589 |
| LOC105668943 | 105668943 | LOC105673321 | 105673321 | spen | 105676238 |
| LOC105668970 | 105668970 | LOC105673333 | 105673333 | ssh | 105677552 |
| LOC105668972 | 105668972 | LOC105673404 | 105673404 | stw | 105668647 |
| LOC105668975 | 105668975 | LOC105673435 | 105673435 | su(w[a]) | 105669558 |
| LOC105669215 | 105669215 | LOC105673486 | 105673486 | syd | 105676167 |
| LOC105669717 | 105669717 | LOC105673498 | 105673498 | tou | 105675153 |
| LOC105669807 | 105669807 | LOC105673529 | 105673529 | trc | 105670855 |
| LOC105669823 | 105669823 | LOC105673632 | 105673632 | trol | 105671774 |
| LOC105670030 | 105670030 | LOC105673654 | 105673654 | trp | 105669468 |
| LOC105670108 | 105670108 | LOC105673737 | 105673737 | tup | 105670267 |
| LOC105670163 | 105670163 | LOC105673760 | 105673760 | twr | 105671606 |
| LOC105670436 | 105670436 | LOC105673816 | 105673816 | unc-104 | 105673707 |
| LOC105670607 | 105670607 | LOC105673909 | 105673909 | unc79 | 105673595 |
| LOC105670882 | 105670882 | LOC105673930 | 105673930 | unpg | 105679002 |
| LOC105670902 | 105670902 | LOC105673989 | 105673989 | upSET | 105669787 |
| LOC105670992 | 105670992 | LOC105673992 | 105673992 | wdb | 105672850 |
| LOC105670997 | 105670997 | LOC105674042 | 105674042 | wg | 105676098 |
| LOC105671033 | 105671033 | LOC105674063 | 105674063 | yem | 105667344 |
| LOC105671038 | 105671038 | LOC105674171 | 105674171 | yki | 105675777 |
| LOC105671243 | 105671243 | LOC105671620 | 105671620 | yps | 105678745 |
| LOC105671415 | 105671415 | LOC105674188 | 105674188 | zfh2 | 105676881 |
| LOC105671611 | 105671611 | LOC105674256 | 105674256 | | |

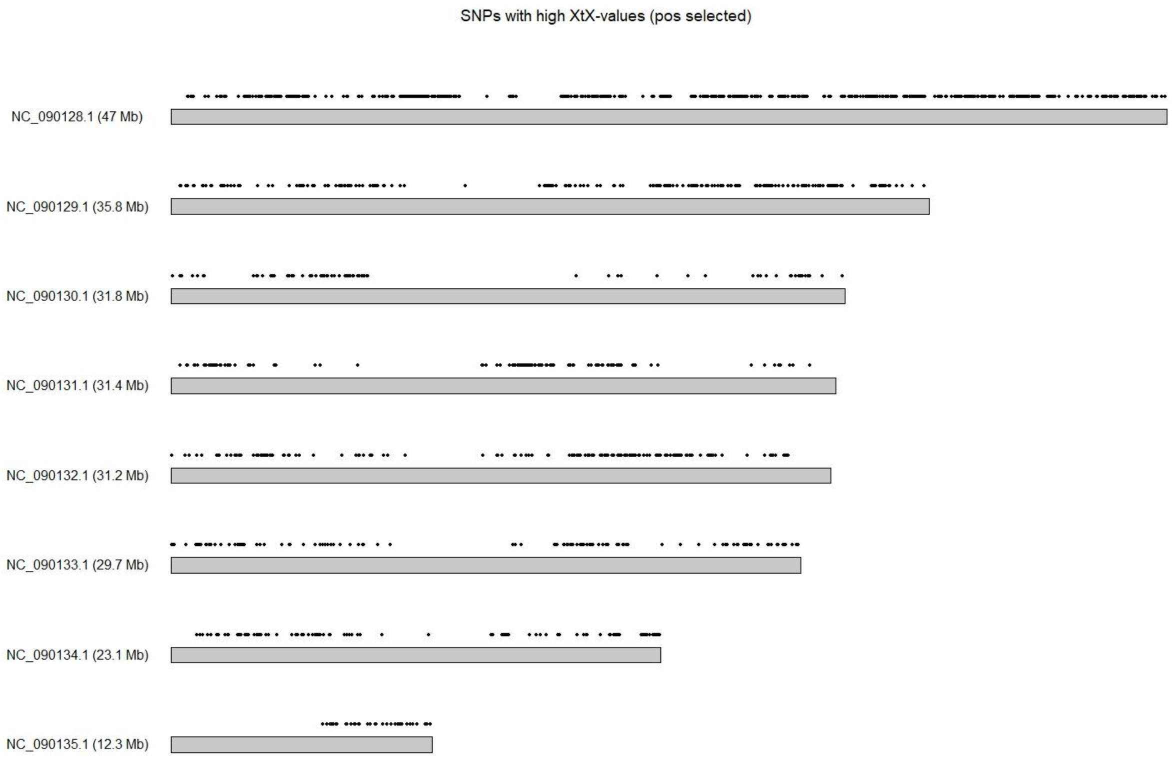


***Figure S9.*** *SNPs with high XtX-values, i.e., candidates for positive selection, were distributed across the genome.*

**Table S10.** Enriched GO terms among positively selected genes.

| Pathway | Fold Enrichment | FDR |
|---|---|---|
| **Molecular Function** | | |
| GO:0015294 Solute:monoatomic cation symporter activity | 8.4365 | 4.9192E-03 |
| GO:0015296 Monoatomic anion:monoatomic cation symporter activity | 8.4365 | 4.9192E-03 |
| GO:0015377 Chloride:monoatomic cation symporter activity | 8.4365 | 4.9192E-03 |
| GO:0004890 Gaba-a receptor activity | 7.9092 | 2.9031E-02 |
| GO:0030215 Semaphorin receptor binding | 7.9092 | 2.9031E-02 |
| GO:0005262 Calcium channel activity | 6.1516 | 6.3334E-04 |
| GO:0015085 Calcium ion transmembrane transporter activity | 5.2728 | 7.0596E-04 |
| GO:0015108 Chloride transmembrane transporter activity | 4.7935 | 2.2504E-02 |
| GO:0015293 Symporter activity | 4.7935 | 2.2504E-02 |
| GO:0099094 Ligand-gated monoatomic cation channel activity | 4.5196 | 1.2873E-02 |
| GO:0015079 Potassium ion transmembrane transporter activity | 4.3637 | 1.5830E-04 |
| GO:0008509 Monoatomic anion transmembrane transporter activity | 4.2183 | 1.8649E-02 |
| GO:0005267 Potassium channel activity | 4.0560 | 1.1987E-03 |
| GO:0015103 Inorganic anion transmembrane transporter activity | 3.9546 | 2.4720E-02 |
| GO:0005261 Monoatomic cation channel activity | 3.4627 | 5.9505E-06 |
| GO:0015291 Secondary active transmembrane transporter activity | 3.3554 | 2.9031E-02 |

| | | |
|---|---|---|
| GO:0046873 Metal ion transmembrane transporter activity | 3.3143 | 1.2484E-05 |
| GO:0005216 Monoatomic ion channel activity | 2.9766 | 2.5591E-07 |
| GO:0015267 Channel activity | 2.9337 | 2.0884E-07 |
| GO:0022803 Passive transmembrane transporter activity | 2.9337 | 2.0884E-07 |
| GO:0015276 Ligand-gated monoatomic ion channel activity | 2.8392 | 3.6247E-03 |
| GO:0022834 Ligand-gated channel activity | 2.8392 | 3.6247E-03 |
| GO:0005230 Extracellular ligand-gated monoatomic ion channel activity | 2.8293 | 1.3131E-02 |
| GO:0022836 Gated channel activity | 2.7829 | 5.7978E-04 |
| GO:0000981 Dna-binding transcription factor activity rna polymerase ii-specific | 2.7179 | 8.3348E-05 |
| GO:0003774 Cytoskeletal motor activity | 2.6364 | 1.0601E-02 |
| GO:0008061 Chitin binding | 2.4813 | 2.4509E-02 |
| GO:0015075 Monoatomic ion transmembrane transporter activity | 2.4421 | 1.1906E-06 |
| GO:0008324 Monoatomic cation transmembrane transporter activity | 2.3892 | 1.7905E-04 |
| GO:0015318 Inorganic molecular entity transmembrane transporter activity | 2.3149 | 5.6363E-04 |
| GO:0003700 Dna-binding transcription factor activity | 2.3085 | 5.8129E-06 |
| GO:0004930 G protein-coupled receptor activity | 2.2533 | 1.2784E-03 |
| GO:0022890 Inorganic cation transmembrane transporter activity | 2.1482 | 5.5146E-03 |
| GO:0140110 Transcription regulator activity | 2.0148 | 7.7496E-05 |
| GO:0022857 Transmembrane transporter activity | 1.7753 | 7.7496E-05 |
| GO:0004672 Protein kinase activity | 1.7497 | 6.3898E-03 |
| GO:0038023 Signaling receptor activity | 1.7143 | 1.8442E-04 |
| GO:0060089 Molecular transducer activity | 1.7143 | 1.8442E-04 |
| GO:0005215 Transporter activity | 1.7121 | 1.5668E-04 |
| GO:0004888 Transmembrane signaling receptor activity | 1.6679 | 6.3334E-04 |
| GO:0003677 Dna binding | 1.5939 | 6.3334E-04 |
| GO:0016773 Phosphotransferase activity alcohol group as acceptor | 1.5681 | 2.6189E-02 |
| GO:0005515 Protein binding | 1.4196 | 2.5591E-07 |
| GO:0005488 Binding | 1.1848 | 1.8894E-05 |
| **Biological Process** | | |
| GO:0048699 Generation of neurons | 5.2728 | 9.0820E-03 |
| GO:0022008 Neurogenesis | 5.2728 | 9.0820E-03 |
| GO:0048731 System development | 4.9213 | 5.8408E-03 |
| GO:0070588 Calcium ion transmembrane transport | 4.9213 | 5.8408E-03 |
| GO:0007399 Nervous system development | 4.9213 | 5.8408E-03 |
| GO:0030182 Neuron differentiation | 4.7935 | 3.6618E-02 |
| GO:0006816 Calcium ion transport | 4.6869 | 3.7674E-03 |
| GO:0048869 Cellular developmental proc. | 4.2183 | 2.9570E-02 |
| GO:0030154 Cell differentiation | 4.2183 | 2.9570E-02 |
| GO:0098609 Cell-cell adhesion | 3.9546 | 4.1198E-02 |
| GO:0007275 Multicellular organism development | 2.7040 | 4.3562E-02 |
| GO:0006813 Potassium ion transport | 2.5778 | 4.2148E-02 |

| | | |
|---|---|---|
| GO:0006811 Monoatomic ion transport | 2.3888 | 9.1340E-06 |
| GO:0034220 Monoatomic ion transmembrane transport | 2.3298 | 8.7653E-03 |
| GO:0030001 Metal ion transport | 2.3069 | 5.8408E-03 |
| GO:0007186 G protein-coupled receptor signaling pathway | 2.0772 | 6.0703E-03 |
| GO:2001141 Reg. of rna biosynthetic proc. | 1.9697 | 7.2530E-06 |
| GO:0006355 Reg. of dna-templated transcription | 1.9697 | 7.2530E-06 |
| GO:0051252 Reg. of rna metabolic proc. | 1.9174 | 1.1787E-05 |
| GO:0019219 Reg. of nucleobase-containing compound metabolic proc. | 1.8862 | 1.9360E-05 |
| GO:0006812 Monoatomic cation transport | 1.8710 | 4.3753E-02 |
| GO:0009889 Reg. of biosynthetic proc. | 1.8056 | 3.7337E-05 |
| GO:0010468 Reg. of gene expression | 1.8056 | 3.7337E-05 |
| GO:0080090 Reg. of primary metabolic proc. | 1.7883 | 5.5125E-05 |
| GO:0031323 Reg. of cellular metabolic proc. | 1.7788 | 5.0506E-05 |
| GO:0016310 Phosphorylation | 1.7499 | 9.1662E-03 |
| GO:0006351 Dna-templated transcription | 1.7451 | 8.3599E-05 |
| GO:0006468 Protein phosphorylation | 1.7419 | 1.1099E-02 |
| GO:0060255 Reg. of macromolecule metabolic proc. | 1.7413 | 8.1860E-05 |
| GO:0055085 Transmembrane transport | 1.7077 | 1.2498E-03 |
| GO:0050794 Reg. of cellular proc. | 1.5957 | 2.0504E-06 |
| GO:0050789 Reg. of biological proc. | 1.5802 | 2.0504E-06 |
| GO:0065007 Biological reg. | 1.5736 | 2.0504E-06 |
| GO:0007165 Signal transduction | 1.5032 | 1.1099E-02 |
| GO:0007154 Cell communication | 1.4809 | 1.3155E-02 |
| GO:0023052 Signaling | 1.4777 | 1.3509E-02 |
| GO:0009987 Cellular proc. | 1.1586 | 9.1662E-03 |

**Table S11.** Genes associated with invasiveness.

| Gene | GeneID | Gene | GeneID | Gene | GeneID |
|---|---|---|---|---|---|
| LOC105671671 | 105671671 | LOC105668476 | 105668476 | LOC136997377 | 136997377 |
| LOC137001523 | 137001523 | LOC105668533 | 105668533 | LOC136997378 | 136997378 |
| LOC105678362 | 105678362 | LOC105668719 | 105668719 | LOC136997379 | 136997379 |
| LOC105668757 | 105668757 | LOC105668732 | 105668732 | LOC136997433 | 136997433 |
| LOC105672731 | 105672731 | LOC105668763 | 105668763 | LOC136997693 | 136997693 |
| LOC105676659 | 105676659 | LOC105668806 | 105668806 | LOC136997746 | 136997746 |
| LOC105676847 | 105676847 | LOC105668813 | 105668813 | LOC136997747 | 136997747 |
| LOC137000480 | 137000480 | LOC105668817 | 105668817 | LOC136997804 | 136997804 |
| LOC137000507 | 137000507 | LOC105668880 | 105668880 | LOC136997910 | 136997910 |
| Pde9 | 105673459 | LOC105669058 | 105669058 | LOC136998114 | 136998114 |
| LOC105672277 | 105672277 | LOC105669338 | 105669338 | LOC136998145 | 136998145 |
| DCX-EMAP | 105680174 | LOC105669933 | 105669933 | LOC136998394 | 136998394 |
| Delta | 105671450 | LOC105669986 | 105669986 | LOC136998396 | 136998396 |

| | | | | | |
|---|---|---|---|---|---|
| Ets96B | 105679624 | LOC105670169 | 105670169 | LOC136998425 | 136998425 |
| LOC105669316 | 105669316 | LOC105670185 | 105670185 | LOC136998814 | 136998814 |
| LOC105670446 | 105670446 | LOC105670217 | 105670217 | LOC136999016 | 136999016 |
| LOC105672074 | 105672074 | LOC105670380 | 105670380 | LOC136999096 | 136999096 |
| LOC105676829 | 105676829 | LOC105670450 | 105670450 | LOC136999721 | 136999721 |
| LOC105679373 | 105679373 | LOC105670460 | 105670460 | LOC136999759 | 136999759 |
| Mctp | 105676442 | LOC105670597 | 105670597 | LOC136999786 | 136999786 |
| Nha1 | 105667833 | LOC105670711 | 105670711 | LOC137000093 | 137000093 |
| Alk | 105679911 | LOC105670731 | 105670731 | LOC137000251 | 137000251 |
| Fas1 | 105674287 | LOC105670742 | 105670742 | LOC137000384 | 137000384 |
| Fas3 | 105671823 | LOC105671055 | 105671055 | LOC137000466 | 137000466 |
| Fhos | 105676030 | LOC105671162 | 105671162 | LOC137000467 | 137000467 |
| Ge-1 | 105671030 | LOC105671223 | 105671223 | LOC137000689 | 137000689 |
| LOC105667797 | 105667797 | LOC105671243 | 105671243 | LOC137001143 | 137001143 |
| LOC105668064 | 105668064 | LOC105671270 | 105671270 | LOC137001147 | 137001147 |
| LOC105668090 | 105668090 | LOC105671360 | 105671360 | LOC137001223 | 137001223 |
| LOC105668191 | 105668191 | LOC105671730 | 105671730 | LOC137001368 | 137001368 |
| LOC105668686 | 105668686 | LOC105671821 | 105671821 | LOC137001629 | 137001629 |
| LOC105668807 | 105668807 | LOC105671854 | 105671854 | LOC137001681 | 137001681 |
| LOC105669315 | 105669315 | LOC105671884 | 105671884 | LOC137001706 | 137001706 |
| LOC105670031 | 105670031 | LOC105671895 | 105671895 | Liprin-beta | 105677523 |
| LOC105670033 | 105670033 | LOC105671897 | 105671897 | MED20 | 105673405 |
| LOC105670144 | 105670144 | LOC105672534 | 105672534 | Mical | 105675006 |
| LOC105670172 | 105670172 | LOC105672654 | 105672654 | Mp | 105674758 |
| LOC105670479 | 105670479 | LOC105672861 | 105672861 | NaPi-III | 105677568 |
| LOC105670483 | 105670483 | LOC105672946 | 105672946 | Nep2 | 105673462 |
| LOC105670882 | 105670882 | LOC105672955 | 105672955 | Nhe2 | 105676318 |
| LOC105671401 | 105671401 | LOC105673404 | 105673404 | Nlg3 | 105674943 |
| LOC105671611 | 105671611 | LOC105673430 | 105673430 | Nmdmc | 105671947 |
| LOC105671620 | 105671620 | LOC105673454 | 105673454 | Octalpha2R | 105675384 |
| LOC105671998 | 105671998 | LOC105673506 | 105673506 | PNKP | 105676740 |
| LOC105672983 | 105672983 | LOC105673842 | 105673842 | PsGEF | 105670183 |
| LOC105673751 | 105673751 | LOC105673872 | 105673872 | Pxn | 105675515 |
| LOC105674520 | 105674520 | LOC105673917 | 105673917 | Rbp6 | 105670561 |
| LOC105674665 | 105674665 | LOC105674017 | 105674017 | Rh50 | 105673446 |
| LOC105674705 | 105674705 | LOC105674045 | 105674045 | Ror | 105673338 |
| LOC105674853 | 105674853 | LOC105674517 | 105674517 | SMC5 | 105676393 |
| LOC105675049 | 105675049 | LOC105674645 | 105674645 | Sap47 | 105671707 |
| LOC105675446 | 105675446 | LOC105674673 | 105674673 | Sar1 | 105671759 |
| LOC105676208 | 105676208 | LOC105674693 | 105674693 | Scr | 105674675 |
| LOC105676460 | 105676460 | LOC105675135 | 105675135 | Set8 | 105673005 |

| | | | | | |
|---|---|---|---|---|---|
| LOC105676680 | 105676680 | LOC105675185 | 105675185 | Sp1 | 105672900 |
| LOC105676831 | 105676831 | LOC105675250 | 105675250 | Spt5 | 105673334 |
| LOC105678148 | 105678148 | LOC105675425 | 105675425 | Syn | 105668648 |
| LOC105678667 | 105678667 | LOC105675447 | 105675447 | Task6 | 105674019 |
| LOC105679342 | 105679342 | LOC105675779 | 105675779 | Ten-m | 105669999 |
| LOC105680116 | 105680116 | LOC105675915 | 105675915 | Timp | 105668698 |
| LOC136997398 | 136997398 | LOC105675932 | 105675932 | Tis11 | 105669901 |
| LOC136997694 | 136997694 | LOC105676222 | 105676222 | Tmtc2 | 105668426 |
| LOC136997990 | 136997990 | LOC105676316 | 105676316 | Toll-6 | 105674053 |
| LOC136998334 | 136998334 | LOC105676320 | 105676320 | Usp1 | 105678972 |
| LOC137000213 | 137000213 | LOC105676363 | 105676363 | Xbp1 | 105677165 |
| LOC137000479 | 137000479 | LOC105676391 | 105676391 | aux | 105675198 |
| LOC137000481 | 137000481 | LOC105676570 | 105676570 | by | 105676351 |
| LOC137000509 | 137000509 | LOC105676573 | 105676573 | cac | 105675747 |
| LOC137001621 | 137001621 | LOC105676576 | 105676576 | cort | 105674680 |
| LanB1 | 105667788 | LOC105676684 | 105676684 | cpx | 105672551 |
| MESR6 | 105673939 | LOC105676690 | 105676690 | crb | 105670455 |
| Nf-YA | 105679218 | LOC105676698 | 105676698 | dlg1 | 105677469 |
| Oseg5 | 105679576 | LOC105676739 | 105676739 | dyl | 105677336 |
| Pgant9 | 105670712 | LOC105676889 | 105676889 | ken | 105677338 |
| Rbfox1 | 105673888 | LOC105676895 | 105676895 | lilli | 105669543 |
| Sema2a | 105670029 | LOC105677049 | 105677049 | mam | 105668331 |
| TfAP-2 | 105673871 | LOC105677558 | 105677558 | msi | 105672415 |
| bru3 | 105671757 | LOC105677686 | 105677686 | mspo | 105672423 |
| corn | 105678811 | LOC105677799 | 105677799 | nvd | 105674672 |
| raskol | 105668128 | LOC105677832 | 105677832 | ple | 105678075 |
| snu | 105672475 | LOC105678008 | 105678008 | pxb | 105670734 |
| Cad87A | 105678923 | LOC105678012 | 105678012 | pygo | 105672898 |
| Chi | 105672223 | LOC105678048 | 105678048 | qin | 105668992 |
| Dll | 105676563 | LOC105678249 | 105678249 | rdgA | 105670288 |
| Dnah3 | 105676990 | LOC105678263 | 105678263 | rtet | 105672557 |
| Fur2 | 105670613 | LOC105678858 | 105678858 | sei | 105669786 |
| HisT | 105671984 | LOC105678982 | 105678982 | sev | 105679157 |
| Imp | 105668782 | LOC105679013 | 105679013 | spri | 105667764 |
| LOC105667419 | 105667419 | LOC105679107 | 105679107 | sprt | 105668800 |
| LOC105667693 | 105667693 | LOC105679626 | 105679626 | stai | 105673759 |
| LOC105667738 | 105667738 | LOC105679779 | 105679779 | unc-13 | 105678225 |
| LOC105667894 | 1056678.94 | LOC105679811 | 105679811 | v | 105680137 |
| LOC105668057 | 105668057 | LOC136996913 | 136996913 | vkg | 105674348 |
| LOC105668385 | 105668385 | LOC136996968 | 136996968 | vvl | 105677135 |
| LOC105668428 | 105668428 | LOC136997375 | 136997375 | zfh2 | 105676881 |

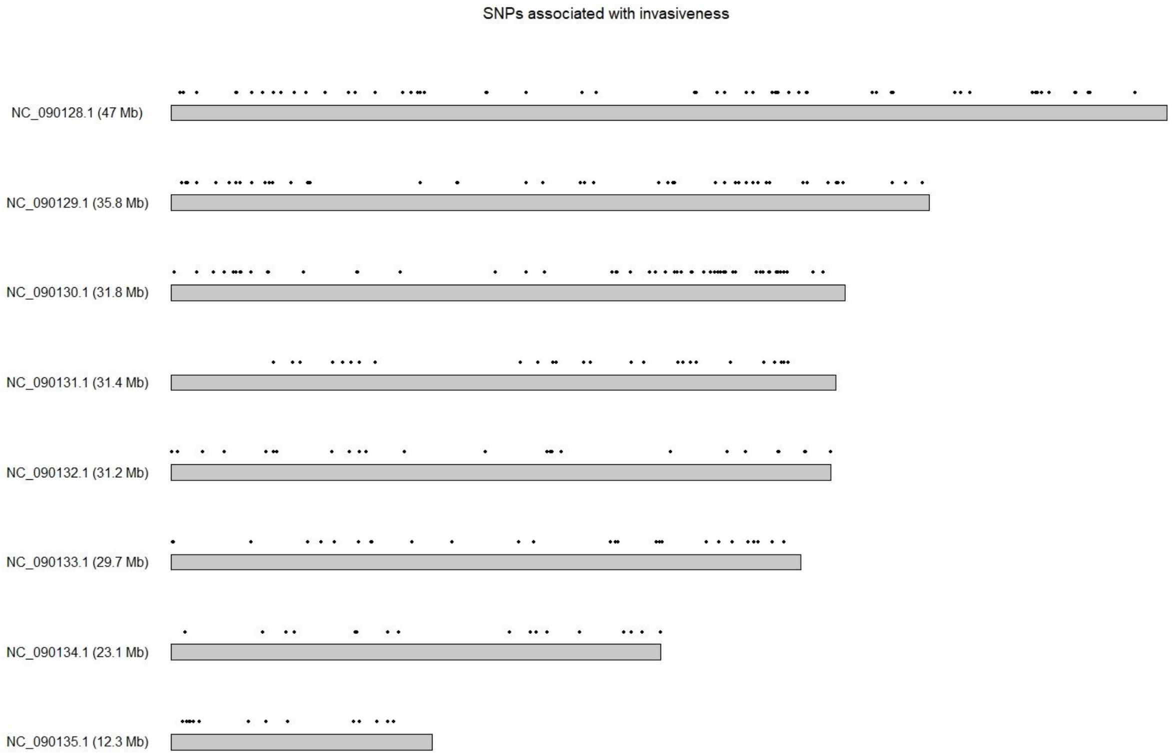


***Figure S12.*** *SNPs with high C2-values, i.e., associated with invasiveness, were distributed across the genome.*

**Table S13.** Enriched GO terms among genes associated with invasiveness.

| **Pathway** | **Fold Enrichment** | **FDR** |
|---|---|---|
| **Molecular Function** | | |
| GO:0003700 Dna-binding transcription factor activity | 3.1881 | 2.7094E-02 |
| GO:0140110 Transcription regulator activity | 3.0057 | 2.7094E-02 |
| **Biological Process** | | |
| GO:0006836 Neurotransmitter transport | 32.8624 | 1.8132E-02 |
| GO:0032886 Reg. of microtubule-based proc. | 32.8624 | 1.8132E-02 |
| GO:0098662 Inorganic cation transmembrane transport | 4.3494 | 3.5669E-02 |
| GO:0098660 Inorganic ion transmembrane transport | 4.2864 | 3.6748E-02 |
| GO:0051252 Reg. of rna metabolic proc. | 2.7159 | 1.3734E-03 |
| GO:2001141 Reg. of rna biosynthetic proc. | 2.6913 | 1.7668E-03 |
| GO:0006355 Reg. of dna-templated transcription | 2.6913 | 1.7668E-03 |
| GO:0009889 Reg. of biosynthetic proc. | 2.6910 | 8.5713E-04 |
| GO:0010468 Reg. of gene expression | 2.6910 | 8.5713E-04 |
| GO:0019219 Reg. of nucleobase-containing compound metabolic proc. | 2.6717 | 1.5628E-03 |

| | | |
|---|---|---|
| GO:0060255 Reg. of macromolecule metabolic proc. | 2.6366 | 8.5713E-04 |
| GO:0031323 Reg. of cellular metabolic proc. | 2.6132 | 1.1238E-03 |
| GO:0080090 Reg. of primary metabolic proc. | 2.5815 | 1.5628E-03 |
| GO:0065007 Biological reg. | 2.4008 | 3.0146E-06 |
| GO:0050789 Reg. of biological proc. | 2.3573 | 5.8292E-06 |
| GO:0050794 Reg. of cellular proc. | 2.2160 | 9.2287E-05 |
| GO:0006351 Dna-templated transcription | 2.2141 | 1.6807E-02 |
| GO:0007154 Cell communication | 2.2025 | 1.0820E-02 |
| GO:0023052 Signaling | 2.1978 | 1.0820E-02 |
| GO:0009987 Cellular proc. | 1.3382 | 1.6807E-02 |

**Table S13.** Enriched GO terms among genes identified with both XtX and C2.

| Pathway | Fold Enrichment | FDR |
|---|---|---|
| **Molecular Function** | | |
| GO:0030215 Semaphorin receptor binding | 68.8846 | 3.3698E-02 |
